# The negative adipogenesis regulator DLK1 is transcriptionally regulated by TIS7 (IFRD1) and translationally by its orthologue SKMc15 (IFRD2)

**DOI:** 10.1101/719922

**Authors:** Ilja Vietor, Domagoj Cikes, Kati Piironen, Theodora Vasakou, David Heimdörfer, Ronald Gstir, Matthias David Erlacher, Ivan Tancevski, Philipp Eller, Egon Demetz, Michael Hess, Volker Kuhn, Gerald Degenhart, Jan Rozman, Martin Klingenspor, Martin Hrabe de Angelis, Taras Valovka, Lukas A. Huber

**Author notes:** Address correspondence to: Ilja Vietor, Institute of Cell Biology, Biocenter, Innsbruck Medical University, Innrain 80-82, A-6020 Innsbruck, Austria. Deceased May 2019 (Volker Kuhn). Conflict of interest: The authors declare that they have no conflict of interest.

## Abstract

Delta-like homolog 1 (DLK1), an inhibitor of adipogenesis, controls the cell fate of adipocyte progenitors. Here we identify two independent regulatory mechanisms, transcriptional and translational, by which TIS7 (IFRD1) and its orthologue SKMc15 (IFRD2) regulate DLK1 levels. Mice deficient in both TIS7 and SKMc15 (dKO) had severely reduced adipose tissue and were resistant to high fat diet-induced obesity. Wnt signaling, a negative regulator of adipocyte differentiation was significantly up regulated in dKO mice. Elevated levels of the Wnt/β-catenin target protein Dlk-1 inhibited the expression of adipogenesis regulators PPARγ and C/EBPα, and fatty acid transporter CD36. Although both, TIS7 and SKMc15, contributed to this phenotype, they utilized two different mechanisms. TIS7 acted by controlling Wnt signaling and thereby transcriptional regulation of Dlk-1. On the other hand, here we provide distinctive experimental evidence that SKMc15 acts as a general translational inhibitor significantly affecting DLK-1 protein levels. Our study provides data describing novel mechanisms of DLK1 regulation in adipocyte differentiation involving TIS7 and SKMc15.

**SYNOPSIS:** 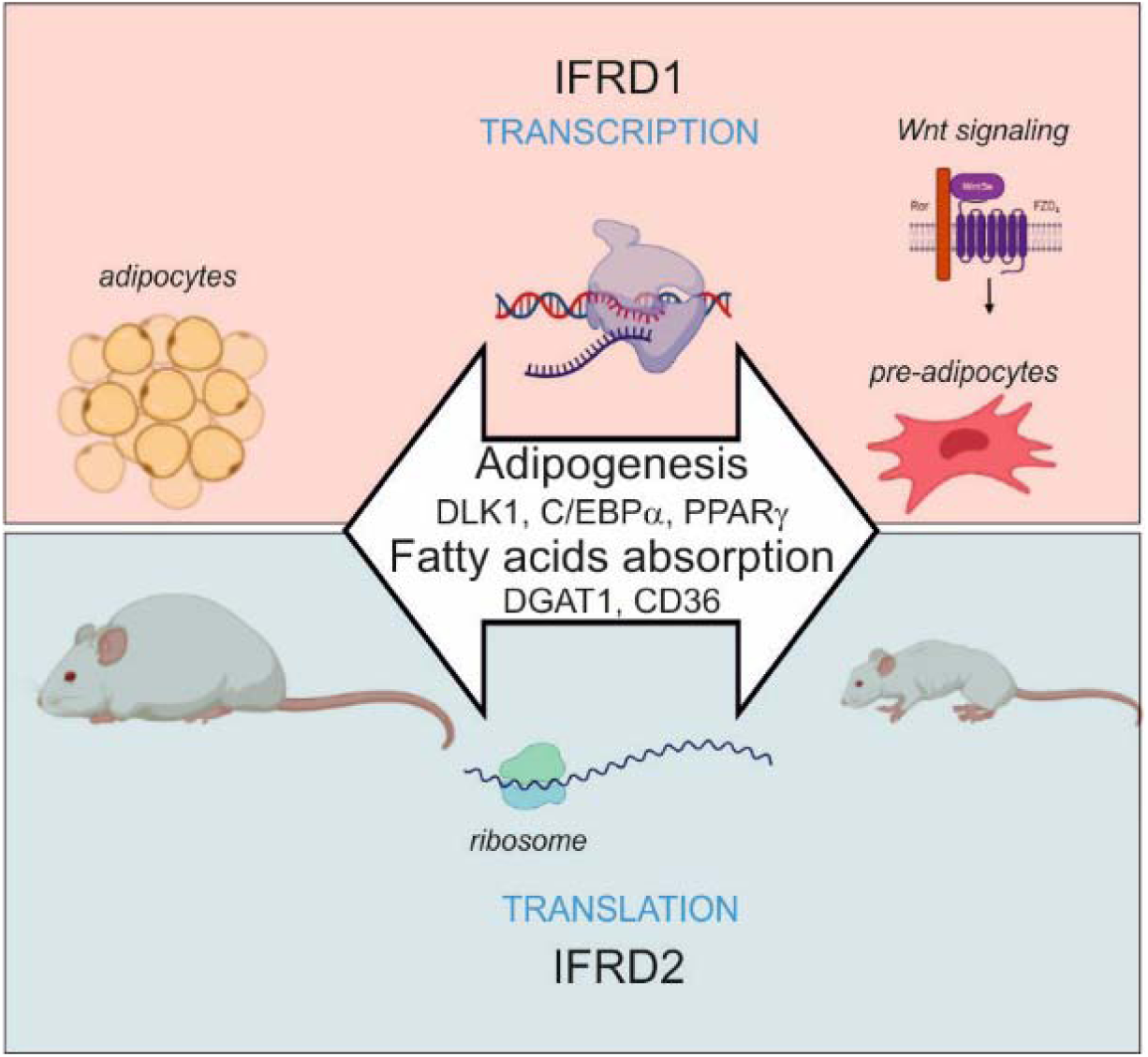

This study uncovered that IFRD1 (TIS7) and its orthologue IFRD2 (SKMc15) are two essential regulators of adipogenesis. These proteins are highly similar on the sequence level, yet they regulate adipocyte differentiation using different but complementary mechanisms. Our main findings are:

- IFRD1 (TIS7) and IFRD2 (SKMc15) knockout mice are resistant against diet-induced obesity
- IFRD1 (TIS7) and IFRD2 (SKMc15) are critical for proper nutritional fat uptake and adipogenesis
- IFRD1 (TIS7) controls adipogenesis via Wnt/β-catenin-dependent transcriptional regulation of adipocyte-specific genes
- IFRD2 (SKMc15) regulates adipocyte-specific genes acting as a novel general translational inhibitor

## Introduction

Adipogenesis is a complex process in which multipotent stem cells are converted into preadipocytes before terminal differentiation into adipocytes (Sarantopoulos, Banyard et al., 2018). These mechanisms involve protein factor regulators, epigenetic factors, and miRNAs. TPA Induced Sequence 7 (TIS7) protein has been shown to be involved in the mainly transcriptional regulation of differentiation processes in various cell types, e.g. neurons (Iacopetti, Barsacchi et al., 1996), enterocytes (Wang, Iordanov et al., 2005), myocytes (Vadivelu, Kurzbauer et al., 2004) and also adipocytes (Nakamura, Hinoi et al., 2013).

Multiple lines of evidence link the regulation of Wnt/β-catenin signaling to the physiological function of TIS7, known in human as Interferon related developmental regulator 1 (IFRD1) (Iezaki, Onishi et al., 2016, Vietor, Kurzbauer et al., 2005). Our data based on experiments with TIS7 knockout mice show a negative effect of TIS7 on Wnt signaling and a positive on adipocyte differentiation (Vietor et al., 2005, Yu, 2010 #3955). TIS7 deficiency leads to a significant up regulation of Wnt/β-catenin transcriptional activity in both primary osteoblasts and in mouse embryonic fibroblasts (MEFs) derived from TIS7 knockout (KO) mice. It was shown that TIS7 is also involved in the control of adipocytes differentiation in mice and was up regulated in both visceral (vWAT) and subcutaneous (sWAT) white adipose tissues (WAT) of genetically obese ob/ob mice (Nakamura et al., 2013). TIS7 transgenic mice have increased total body adiposity and decreased lean mass compared with wild type (WT) littermates (Wang et al., 2005). On HFD, TIS7 transgenic mice exhibit a more rapid and proportionately greater gain in body weight with persistently elevated total body adiposity. Enhanced triglyceride (TG) absorption in the gut of TIS7 transgenic mice (Wang et al., 2005) indicated that TIS7 expressed in the gut epithelium has direct effects on fat absorption in enterocytes. We have shown that as a result of impaired intestinal lipid absorption TIS7 KO mice displayed lower body adiposity (Yu, Jiang et al., 2010). Compared with WT littermates, TIS7 KO mice do not gain weight when chronically fed an HFD and TIS7 deletion results in delayed lipid absorption and altered intestinal and hepatic lipid trafficking, with reduced intestinal TG, cholesterol, and free fatty acid mucosal levels in the jejunum (Garcia, Wakeman et al., 2014). TIS7 protein functions as a transcriptional co-regulator (Micheli, Leonardi et al., 2005) due to its interaction with protein complexes containing either histone deacetylases (HDAC) (Park, Horie et al., 2017, Vadivelu et al., 2004, Vietor, Vadivelu et al., 2002, Wick, Schleiffer et al., 2004) or protein methyl transferases, in particular PRMT5 (Lammirato, Patsch et al., 2016). The analysis of adipocyte differentiation in preadipocytic 3T3-L1 cells suggested an involvement of TIS7 in the regulation of adipogenesis in the Wnt/β-catenin signaling context (Nakamura et al., 2013).

SKMc15, also known as Interferon related developmental regulator 2 (IFRD2), a second member of TIS7 gene family, is highly conserved in different species (Latif, Duh et al., 1997). Mouse TIS7 and SKMc15 are highly homologous, with a remarkable identity at both the cDNA and amino acid levels (58% and 88%, respectively). However, there was so far no information about the physiological function and mechanisms of action of SKMc15 and its possible involvement in differentiation of various tissues. Recently, cryo-electronmicroscopy (cryo-EM) analyses of inactive ribosomes identified IFRD2 (SKMc15) as a novel ribosome-binding protein inhibiting translation to regulate gene expression (Brown, Baird et al., 2018). The physiological function of SKMc15 matching the mechanism based on the above mentioned cryo-EM data was so far not shown and we postulate it could be involved in adipogenesis, since a significant reduction of whole protein synthesis was previously shown as a major regulatory event during early adipogenic differentiation (Marcon, Holetz et al., 2017).

Adipogenesis occurs late in embryonic development and in postnatal periods. Adipogenic transcription factors CCAAT/enhancer binding protein α (C/EBPα) and peroxisome proliferator activated receptor γ (PPARγ) play critical roles in adipogenesis and in the induction of adipocyte markers (Farmer, 2006). PPARγ is the major downstream target of Delta-like protein 1 (DLK-1). It is inactivated by the induction of the MEK/ERK pathway leading to its phosphorylation and proteolytic degradation (Wang & Sul, 2009). DLK-1, also known as Pref-1 (preadipocyte factor 1) activates the MEK/ERK pathway to inhibit adipocyte differentiation (Kim, Kim et al., 2007). C/EBPα is highly expressed in mature adipocytes and can bind DNA together with PPARγ to a variety of respective target genes (Lefterova, Zhang et al., 2008). Besides that, PPARγ binding to C/EBPα gene induces its transcription, thereby creating a positive feedback loop (Lowell, 1999). Both proteins have synergistic effects on the differentiation of adipocytes that requires a balanced expression of both C/EBPα and PPARγ.

Wnt/β-catenin signaling is one of the extracellular signaling pathways specifically affecting adipogenesis (Li, Luo et al., 2008, Ross, Hemati et al., 2000, van Tienen, Laeremans et al., 2009) by maintaining preadipocytes in an undifferentiated state through inhibition of C/EBPα and PPARγ (Tontonoz & Spiegelman, 2008). PPAR-γ and Wnt/β-catenin pathways are regarded as master mediators of adipogenesis (Xu, Wang et al., 2016). Wnt signaling is a progenitor fate determinator and negatively regulates preadipocyte proliferation through DLK-1 (Mortensen, Jensen et al., 2012). Mice overexpressing DLK-1 are resistant to HFD-induced obesity, whereas DLK-1 KO mice have accelerated adiposity (Moon, Smas et al., 2002). DLK-1 transgenic mice show reduced expression of genes controlling lipid import (CD36) and synthesis (Srebp1c, Pparγ) (Barclay, Nelson et al., 2011). DLK-1 expression coincides with altered recruitment of PRMT5 and β-catenin to the DLK-1 promoter (Paul, Sardet et al., 2015). PRMT5 acts as a co-activator of adipogenic gene expression and differentiation (LeBlanc, Konda et al., 2012). SRY (sex determining region Y)-box 9 (Sox9), a transcription factor expressed in preadipocytes, is down regulated preceding adipocyte differentiation. DLK-1 prevents down regulation of Sox9 by activating ERK, resulting in inhibition of adipogenesis (Sul, 2009). The PRMT5-and histone-associated protein COPR5, affects PRMT5 functions related to cell differentiation (Paul, Sardet et al., 2012). Adipogenic conversion is delayed in MEFs derived from COPR5 KO mice and WAT of COPR5 KO mice is reduced when compared to control mice. DLK-1 expression is up regulated in COPR5 KO cells (Paul et al., 2015).

Here we show the involvement of TIS7 and SKMc15 in the process of adipocyte differentiation. dKO mice had strongly decreased amounts of the body fat when fed with even regular, chow diet and were resistant against the high fat diet induced obesity. We found two independent molecular mechanisms through which TIS7 and SKMc15 fulfill this function. The fact that these two genes use independent mechanisms of action supported by the observation that whole body deficiency of both genes led to a stronger phenotype when compared to single knockouts of TIS7 or SKMc15. TIS7 regulates the Wnt signaling pathway activity and restricts DLK-1 protein levels, thereby allowing adipocyte differentiation. In contrast, SKMc15 KO did not affect Wnt signaling, but as we show here, cells lacking SKMc15 have significantly up regulated translational activity. In addition, we identified strongly enriched Dlk-1 mRNA concentrations specifically in polyribosomes isolated from SKMc15 knockout MEFs when compared to the WT MEFs. This was true also for dKO, but not for the single TIS7 knockout cells. The ablation of both TIS7 and SKMc15 genes significantly affected the expression of genes essential for adipocyte differentiation and function. Since dKO mice render a substantially leaner phenotype on chow diet, even without any challenge by HFD-induction, we propose that TIS7 and SKMc15 represent novel players in the process of physiological adipocyte differentiation.

## Results

### Mice lacking TIS7 and SKMc15 genes have lower body mass, less fat and are resistant against HFD-induced obesity

In order to clarify whether both, TIS7 and SKMc15 are involved in the regulation of the adipocyte differentiation and if they act through same or different mechanisms, we have generated mice lacking both genes by crossing TIS7 with SKMc15 single KO mice. dKO pups were viable, and adult male and female mice were fertile. At birth, body weights of dKO and WT mice were similar. Nevertheless, already during weaning, both the male and female dKO mice failed to gain weight when compared to their WT littermates and this persisted in the following weeks when the mice were fed regular diet (RD; chow diet) containing 11 % kcal of fat. At 10 weeks of age, dKO mice displayed 30 % and later up to 44.9 % lower body weight compared with WT mice (Fig 1A).

**Figure 1.**
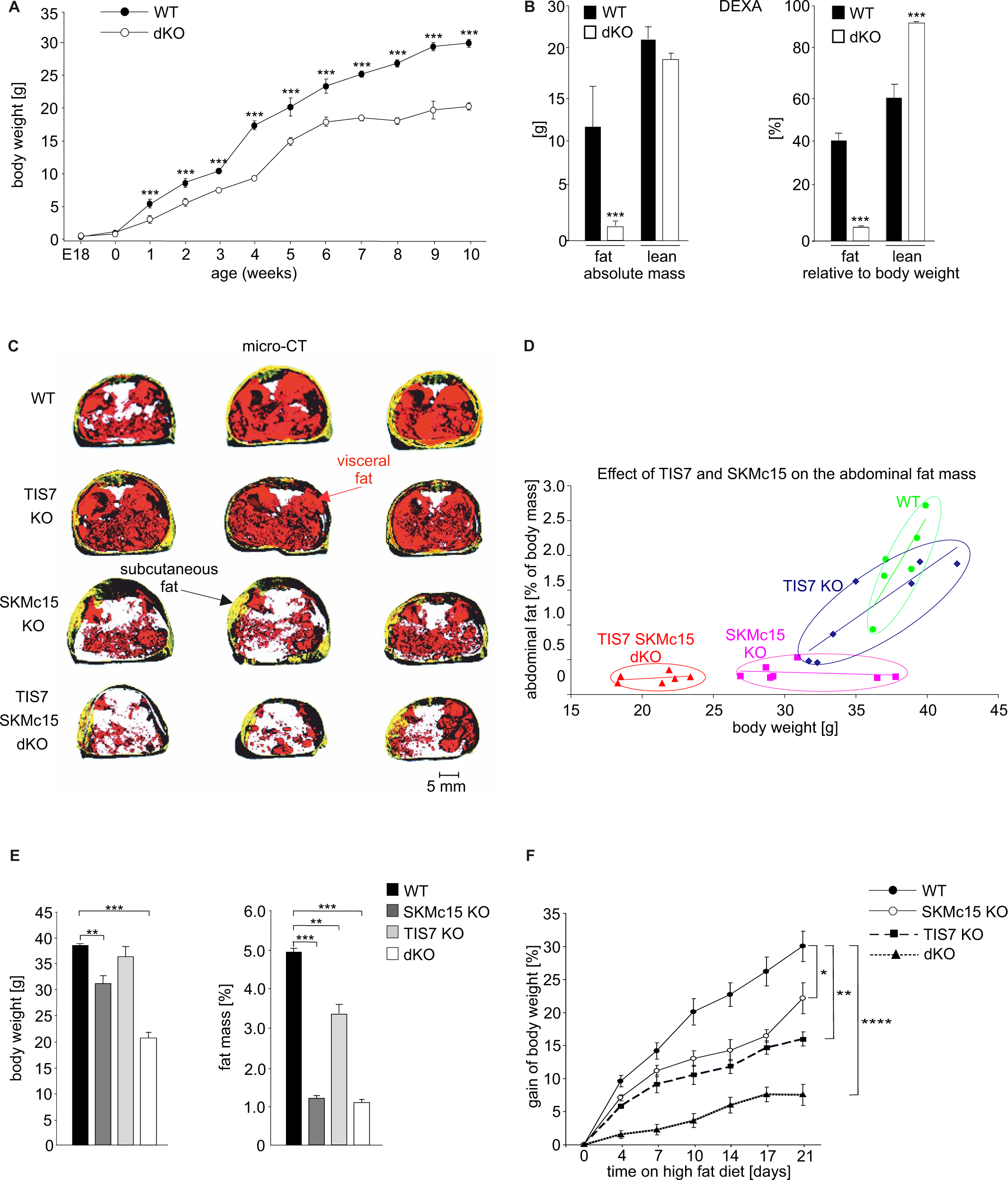
dKO mice display postnatal growth retardation, less body fat and are resistant against HFD-induced obesity. A) Growth curves of WT and dKO mice on chow diet (n fluctuate depending on the time point of measurement). B) DEXA measurements of WT and dKO mice. Left panel depicts the absolute values of fat and lean mass per animal and the right panel represents values normalized to the total body weight of animals (n=15). C) Micro-computed tomography measurements identified a lack of abdominal fat in single and dKO mice. 3-dimensional reconstitution of images of the abdominal fat mass distribution in WT and KO mice. Yellow color represents subcutaneous and red visceral fat mass. Black compartments are shadows resulting as part of the lightning model for the 3D volume rendering. D) Mass contribution [%] of the abdominal fat in correlation to the total body weight [g]. A linear regression is overlaid for each group individually. The regression results for the WT group are y=0,342x-11,13 with a R^2^ of 0,74; for the TIS7 KO group y=0,143x-3,91 with a R^2^ of 0,82; for the SKMc15 KO y=0,005x+0,48 with a R^2^ of 0,04 and for the dKO y=0,011x+0,01 with a R^2^ of 0,1. ANCOVA analysis for the fat mass as a percentage of the body weigth as covariant was performed. E) Effect of single SKMc15 or TIS7 knockout and TIS7 SKMc15 double knockout on total body weight in [g] and body fat amount normalized to the body weight in [%]. F) TIS7 and SKMc15 knockout reduced the gain of body weight of mice fed with HFD. 7 weeks old male WT and dKO mice were caged individually and maintained up to 21 days on HFD. Data shown are mean ± STD, n≥9 per genotype. Data were analyzed applying one-way ANOVA with Holm-Šidák’s multiple comparisons test. *P<0.05, **P<0.01, **** P<0.0001.

Based on Dual Energy X-ray Absorptiometry (DEXA) measurement, 6-month-old WT mice had substantially higher amounts of fat than their dKO littermates (Fig 1B, left panel). The effect of TIS7 and SKmc15 double knockout was even more pronounced when the total fat and lean mass values were normalized to the body weight since the dKO mice were smaller than their WT counterparts were. The percentage of fat was in the WT mice 37.7 ± 4 % vs. 6 ± 3 % of the total body mass in dKO animals. Furthermore, the percentage of lean tissue mass in WT animals was lower than in dKO animals (60 ± 4.6 % vs. 92 ± 3.14 %; Fig 1B, right panel). Nevertheless, the dKO mice were not significantly smaller, since there was only a minor difference in body length, including the tail differed between WT and dKO mice (Fig EV1A). Next, we analyzed the contribution of TIS7 and SKMc15 to the whole-body fat content of mice. Three-dimensional reconstruction of images based on micro-computed tomography (micro-CT) of sex/age-matched adult mice disclosed that both TIS7 and SKMc15 single KO mice already had less abdominal fat than WT controls and that the dKO of TIS7 and SKMc15 genes caused the strongest decrease in abdominal fat content and size (Fig 1C). Quantitative analyses of micro-CT measurements showed that TIS7 deficiency caused a substantial lack of the abdominal fat tissues (P=0.002 when compared to WT mice, Fig 1D). Whereas TIS7 KO mice had less fat mass but were not significantly lighter than their WT littermates (Fig 1E), SKMc15 KO were lighter (P=0.007) and leaner (P=0.0001) and dKO mice were both significantly lighter (P = 0.0001) and had significantly less abdominal fat (P=0.006) than the WT mice (Fig 1D,E).

The indirect calorimetry trial with sex-and age-matched WT and dKO animals where did not identify any significant difference in respiratory exchange ratios (RER = VCO2/VO2) of WT and dKO mice (Fig EV1B). However, dKO mice showed substantially reduced body weight, mainly because of lacking fat, despite identical food intake, activity and no major differences in several metabolic parameters. To investigate potential links between TIS7, SKMc15 and obesity, we studied the response of dKO mice to HFD. At 2 months of age, male mice were housed individually and fed with HFD for 21 days. Food intake was measured every second day and body weight was measured every fourth day. Feces were collected every second day to analyze the composition of excreted lipids and blood samples were collected after the third week of HFD feeding to measure the concentrations of hepatic and lipoprotein lipases, respectively. Already within the first week of HFD feeding, WT mice gained more weight than the dKO mice (Fig 1F), although there were no obvious differences in food consumption (Suppl fig 1A) or in levels of lipolytic enzymes (Suppl fig 1B,C). These differences in body weight gain continued to increase during the second and third week, at which time the body weight of WT mice increased additionally for 30% (30.0 ± 2.3%) when compared with the beginning of the HFD feeding period. In contrast, the weight of dKO animals increased only slightly (7.6 ± 1.6%) (Fig 1F). Both genes, TIS7 and SKMc15 contributed to this phenotype and we could see a stronger effect following their deletion (Fig 1F).

### Adipocyte differentiation in dKO mice is inhibited due to up regulated DLK-1 levels

A possible explanation of the lean phenotype of dKO mice was that TIS7 and SKMc15 regulate adipocyte differentiation. Primary MEFs derived from totipotent cells of early mouse mammalian embryos are capable of differentiating into adipocytes and they are versatile models to study adipogenesis as well as mechanisms related to obesity such as genes, transcription factors and signaling pathways implicated in the adipogenesis process (Ruiz-Ojeda, Ruperez et al., 2016). To test whether TIS7 and SKMc15 are required for adipogenesis we treated MEFs derived from WT, SKMc15 and TIS7 single and dKO mice using an established adipocyte differentiation protocol (Wang, Sun et al., 2015). Expression levels of both TIS7 and SKMc15 mRNA increased during the adipocyte differentiation of WT MEFs. TIS7 expression reached maximum levels representing 5.7-fold increase compared to proliferating WT MEFs on day 3 (Suppl fig 1D) and SKMc15 reached on day 5 the maximum of 2.5-fold expression levels of proliferating MEFs (Suppl fig 1E). These data suggested that both proteins play a regulatory role in adipogenesis, however differ in their mechanisms and timing. Eight days after initiation of adipocyte differentiation a remarkable reduction of adipocyte differentiation potential in SKMc15, TIS7 KO MEFs and dKO stromal vascular fraction (SVF) cells isolated from inguinal WAT was observed, as characterized by the formation of lipid droplets stained by oil red O (Fig 2A). Quantification of this staining revealed that fat vacuole formation in cells derived from SKMc15, TIS7 and dKO mice represented 23 %, 48 % and 12 % of the WT cells, respectively (Fig 2B). Stable ectopic expression of SKMc15 significantly increased adipocyte differentiation in both, single and double TIS7 and SKMc15 knockout MEF cell lines (Fig EV1C,D and EV2A). Ectopic expression of TIS7 significantly induced the adipocyte differentiation in TIS7 single knockout MEFs (Fig EV1C). These data indicated that both, TIS7 and SKMc15 were critical for adipocyte differentiation and that the defect in adipogenesis could be responsible for the resistance of dKO mice to HFD-induced obesity.

**Figure 2.**
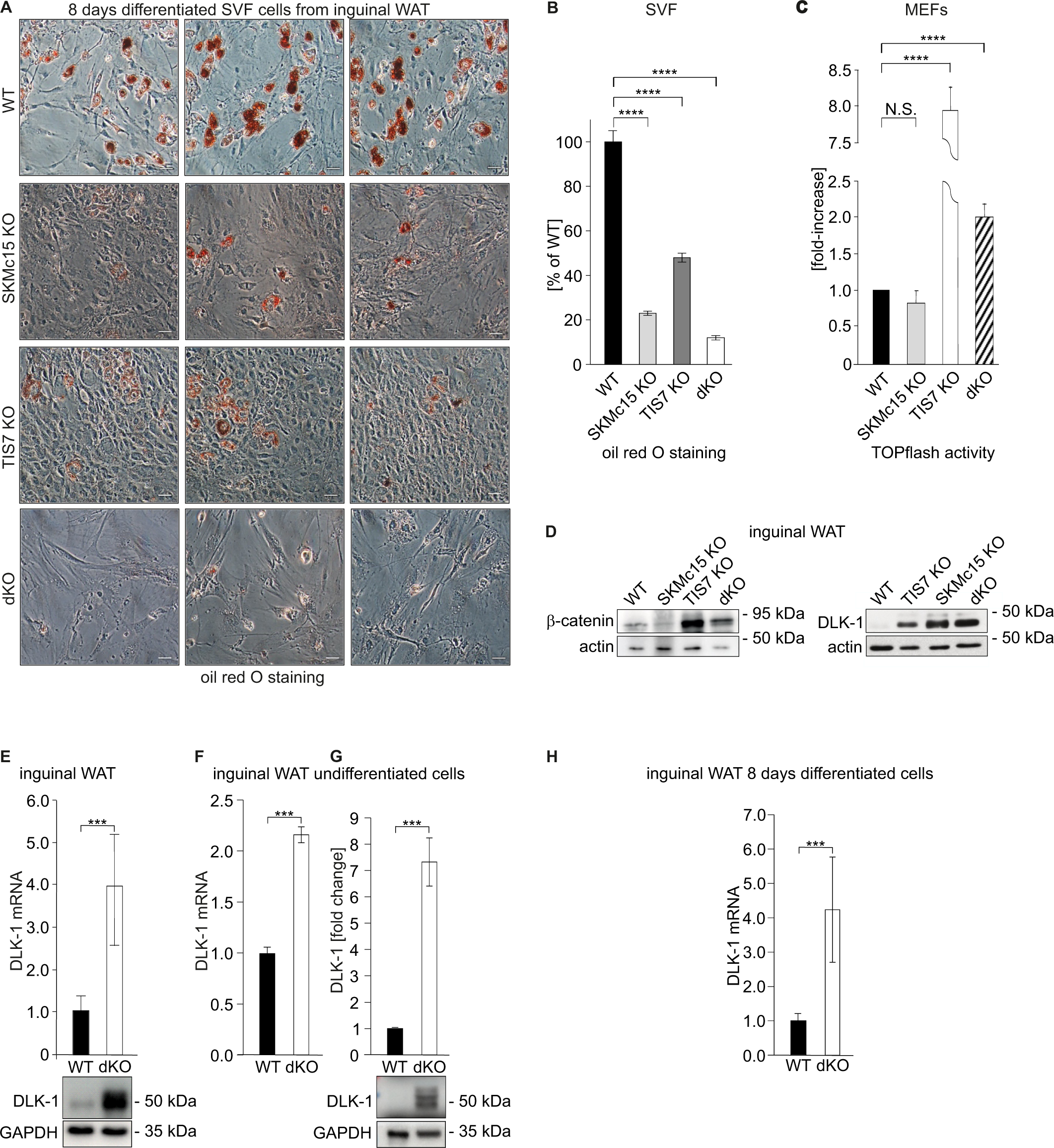
Adipocyte differentiation, DLK-1 levels and Wnt signaling pathway are significantly affected in dKO mice. A) Representative oil red O staining of inguinal SVF cells following 8 days of the adipocyte differentiation protocol. Scale bar is 20µm. B) Quantification of oil red O staining from three independent adipocyte differentiation experiments shown in panel A. Ordinary one-way ANOVA **** P<0.0001. C) Wnt signaling activity was measured in MEFs transiently co-transfected with the Tcf reporter plasmid pTOPflash and β-galactosidase expression vectors. Transfection efficiency was normalized using the β-galactosidase values. The experiment was repeated three times. Data shown are mean ± STD. Ordinary one-way ANOVA Holm-Šídák’s multiple comparisons test **** P<0.0001. D) Left top panel: Western blot analysis of β-catenin amounts in inguinal WAT samples. Representative Western blot from 3 biological repetitions is shown. Right top panel: DLK-1 is significantly up regulated both in TIS7, SKMc15 single KO and in dKO mice. Bottom panels: actin Western blots were used for normalization of sample loading. E) DLK-1 mRNA expression measured by qPCR in inguinal WAT tissue samples (top). Values were normalized on GAPDH, n=3. Error bars indicate standard deviations. A representative Western blot image detecting DLK-1 and GAPDH proteins (bottom). F) Dlk-1 mRNA expression detected in undifferentiated cells isolated from inguinal WAT SVF cells. Normalized on GAPDH. Error bars indicate standard deviations, ***P<0.001. G) DLK-1 protein Western blot analysis in undifferentiated cells isolated from inguinal WAT SVF cells. Mean of 3 biological repeats; insert is one representative Western blot. ***P<0.001. H) Dlk-1 mRNA expression detected in 8 days adipocyte differentiated cells isolated from inguinal WAT SVF cells. Normalized on GAPDH. Error bars indicate standard deviations, ***P<0.001.

Wnt/β-catenin signaling is an important regulatory pathway for adipocyte differentiation (Prestwich & Macdougald, 2007). Consistently, others and we have shown that TIS7 deficiency causes up regulation of Wnt/β-catenin transcriptional activity. Therefore, we have assessed Wnt signaling activity in MEFs by measuring transcriptional activity using TOPflash, TCF-binding luciferase reporter assays. As shown in Fig 2C, Wnt signaling activity, when compared to the WT MEFs, was not regulated in SKMc15 KO, but highly significantly (P<0.0001) up regulated in TIS7 KO and also in dKO MEFs. Supporting these findings, Western blot analyses also identified increased β-catenin protein levels in inguinal WAT of TIS7 KO and in dKO mice (P<0.0001), but not in SKMc15 KO mice (Fig 2D, left top panel) suggesting the involvement of TIS7 but not SKMc15 in the Wnt signaling pathway regulation.

Next, we analyzed the expression levels of DLK-1, a negative regulator of adipogenesis and at the same time known target of Wnt signaling (Paul et al., 2015). DLK-1 protein was significantly up regulated in inguinal WAT isolated from TIS7 and SKMc15 KO as well as from dKO mice (Fig 2D, right top panel). A significant (P<0.001) up regulation of DLK-1 mRNA and protein levels in dKO inguinal WAT samples revealed the qPCR and confirmed Western blot analyses (Fig 2E). Up regulation of DLK-1 in dKO mice both on RNA and protein results were confirmed in undifferentiated SVF cells isolated from inguinal fat (Fig 2F,G). DLK-1 mRNA levels were even stronger up regulated following the 8 days differentiation protocol of SVF cells (Fig 2H). Moreover, whereas in WT MEFs the expression of DLK-1 was strongly up regulated only during the first day of adipocyte differentiation and then over the next days declined to basal levels, in dKO MEFs DLK-1 was up regulated (P<0.001) throughout the entire 8 days of differentiation (Fig 3A). This result complemented protein analyses of lysates from 8 day differentiated adipocytes (Fig EV2B middle panel). A rescue experiment confirmed that DLK-1 expression was TIS7-and SKMc15-dependent. DLK-1 mRNA levels were analyzed by RT-qPCR in dKO MEFs stably expressing TIS7 and/or SKMc15. Ectopic expression of TIS7, SKMc15 and mainly their combination significantly (P<0.001) down regulated DLK-1 RNA and protein levels (Fig 3B). Accordingly, these experiments documented that TIS7 and/or SKMc15 were involved in the regulation of DLK-1 expression, but the molecular mechanism remained unclear.

**Figure 3.**
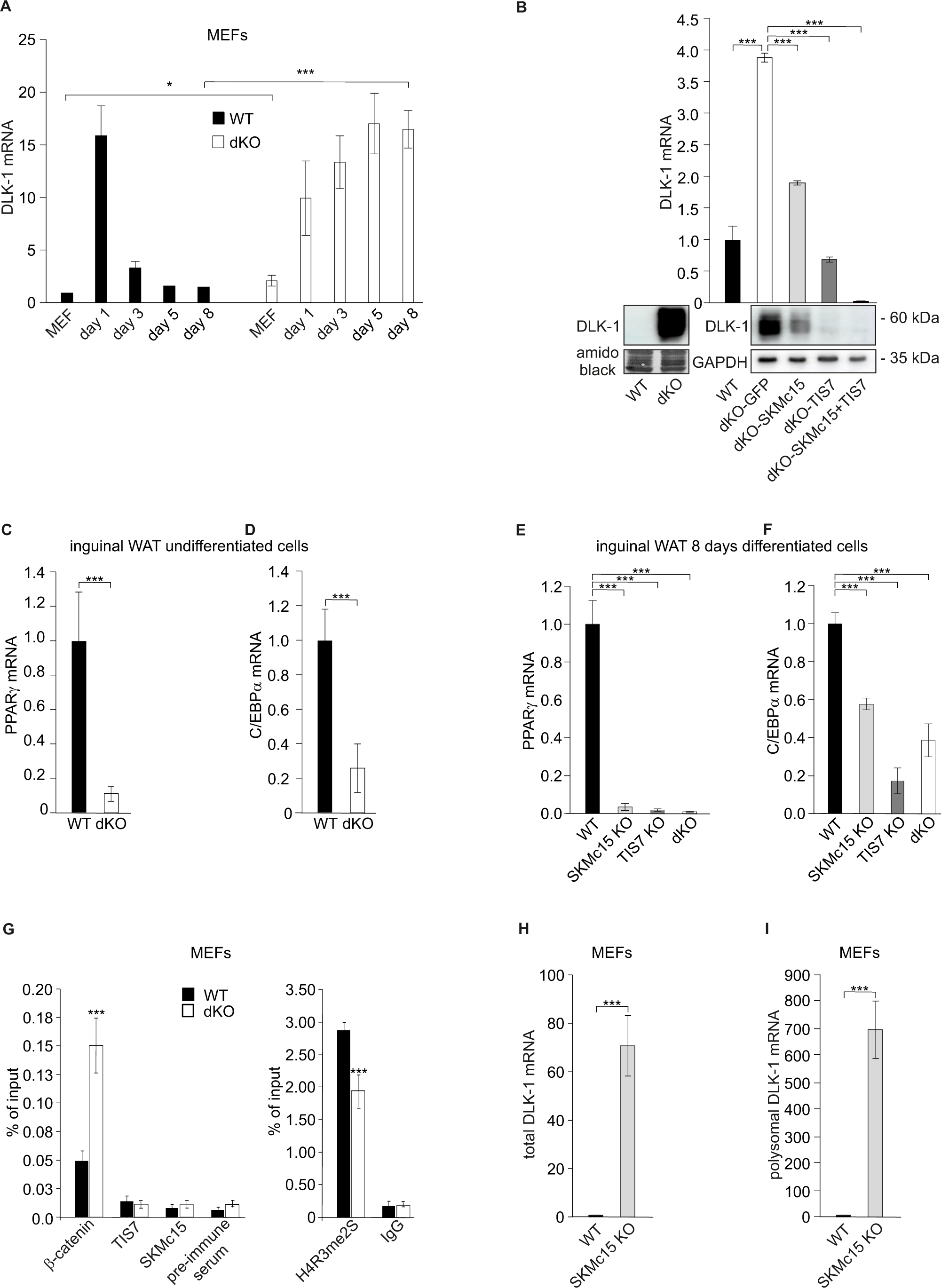
TIS7 and SKMc15 regulate Dlk-1 and thereby affect adipocyte regulators expression. A) DLK-1 mRNA expression measured in MEFs treated with the adipocyte differentiation cocktail for given times. WT MEFs values were set as 1. *P<0.05, ***P<0.001. B) DLK-1 mRNA levels (top) and protein levels (bottom) in dKO MEFs were down regulated following ectopic co-expression of TIS7 and SKMc15. mRNA expression levels were analyzed by RT qPCR in stably transfected cells following 8 days adipocyte protocol differentiation. WT MEFs values were set as 1, n=3; Error bars indicate standard deviations. *** P<0.001 C) PPARγ mRNA expression detected in undifferentiated cells isolated from inguinal WAT SVF cells. Normalized on GAPDH. Error bars indicate standard deviations, ***P<0.001. D) C/EBPα mRNA expression detected in undifferentiated cells isolated from inguinal WAT SVF cells. Normalized on GAPDH. Error bars indicate standard deviations, ***P<0.001. E) PPARγ mRNA expression detected in 8 days adipocyte differentiated cells isolated from inguinal WAT SVF cells. Normalized on GAPDH. Error bars indicate standard deviations, ***P<0.001. F) C/EBPα mRNA expression detected in 8 days adipocyte differentiated cells isolated from inguinal WAT SVF cells. Normalized on GAPDH. Error bars indicate standard deviations, ***P<0.001. G) Recruitment of indicated proteins to regulatory regions of the Dlk-1 promoter in WT and KO MEFs was analyzed by ChIP at day 8 of the adipocyte differentiation. Values are expressed as the percentage of immunoprecipitated chromatin relative to input and are the mean of triplicates. ChIP analysis identified increased specific β-catenin binding to its DLK-1 regulatory element in dKO samples. Pre-immune serum and IgG were used as background controls. n=3, Data shown are mean ± STD, Student’s t-test ***P<0.001. H) Dlk-1 mRNA expression detected in undifferentiated MEF cells. Normalized on GAPDH. Error bars indicate standard deviations, ***P<0.001. I) Real-time qPCR detection of Dlk-1 RNA in polyribosomes. Normalized on GAPDH. Error bars indicate standard deviations, ***P<0.001.

The adipocyte differentiation deficiency of dKO inguinal SVF cells suggested that C/EBPα and PPARγ might also be regulated through TIS7 and/or SKMc15. It was previously shown that elevated levels of the cleaved ectodomain of DLK1 have been correlated with reduced expression of PPARγ (Lee, Villena et al., 2003). Therefore, we analyzed the differences in PPARγ expression between WT and dKO inguinal SVF cells. While PPARγ and C/EBPα mRNA levels were strongly induced in undifferentiated WT, these were barely detectable in dKO SVF cells (Fig 3C,D). Similarly, we identified significantly decreased PPARγ and C/EBPα mRNA levels in TIS7, SKMc15 and dKO inguinal SVF cells following 8 days adipocyte differentiation protocol (Fig 3E,F).

Our earlier study revealed that TIS7 binds directly to DNA and via interaction with PRMT5 regulates gene expression (Lammirato et al., 2016). Therefore, we performed chromatin immunoprecipitation (ChIP) experiments where we studied binding of TIS7 and SKMc15 as well as of the transcription factor β-catenin and symmetrically dimethylated histone H4 at arginine residue 3 (H4R3me2s) (Paul et al., 2015) to the regulatory elements of the DLK-1 gene. In dKO MEFs treated 8 days with the adipocyte differentiation cocktail we found increased β-catenin binding to the β-catenin/TCF binding site 2 of the Dlk-1 regulatory element when compared to WT MEFs (Fig 3G). On the other hand, binding of H4R3me2s to the same Dlk-1 regulatory element was significantly reduced (P<0.001). We could not identify any direct binding of TIS7 or SKMc15 proteins to two different DLK-1 regulatory elements (Dlk-1 region A and β-cat/TCFbs2), neither in WT nor in dKO MEFs, suggesting rather an epigenetic regulation than *via* their direct binding to DLK-1 regulatory elements. We concluded that TIS7 and SKMc15 are required to restrain the DLK-1 levels, through the Wnt/β-catenin signaling pathway and yet another so far unknown mechanism.

After finding significantly increased DLK-1 protein levels in inguinal WAT of SKMc15 single knockout mice (Fig 2D) we measured DLK-1 expression in MEFs generated from these mice. These were significantly increased (>70-fold) when compared to WT MEFs (Fig 3H). In a search for a SKMc15-specific regulatory mechanism of Dlk-1 levels we focused on the translational regulation. It was previously shown that general reduction of protein synthesis and downregulation of the expression and translational efficiency of ribosomal proteins are events crucial for the regulation of adipocyte differentiation (Marcon et al., 2017). Initially, we have measured specifically polyribosome-bound DLK-1 RNA in SKMc15 knockout MEFs. Our analysis identified significantly (P<8,26104 E-07) higher DLK-1 mRNA levels in polyribosome RNA fraction of SKMc15 KO when compared to wild type MEFs (Fig 3I). Interestingly, IFRD2 (SKMc15) was recently identified as a novel specific factor capable of translationally inactivating ribosomes (Brown et al., 2018). Because SKMc15 may play a crucial role in the adipogenesis regulation we tested in our following experiment the effect of SKMc15 knockout on the general translational efficiency of WT, TIS7, SKMc15 single and double KO MEFs. Cells were incubated 30 minutes in the absence of methionine and cysteine, followed by one hour in the presence of ^35^S-methionine. As shown in Fig 4A, there was a significant increase in the general translational activity of MEFs lacking SKMc15 alone or both TIS7 and SKMc15, but not in TIS7 single knockout cells. Therefore, we concluded that SKMc15 alone, but not TIS7 inhibits the general translational activity necessary for the induction of adipogenic differentiation, also via the translational regulation of Dlk-1. This finding supported the result shown in Fig 2D where the DLK-1 protein levels were significantly induced in inguinal WAT samples isolated from SKMc15 single knockout mice.

**Figure 4.**
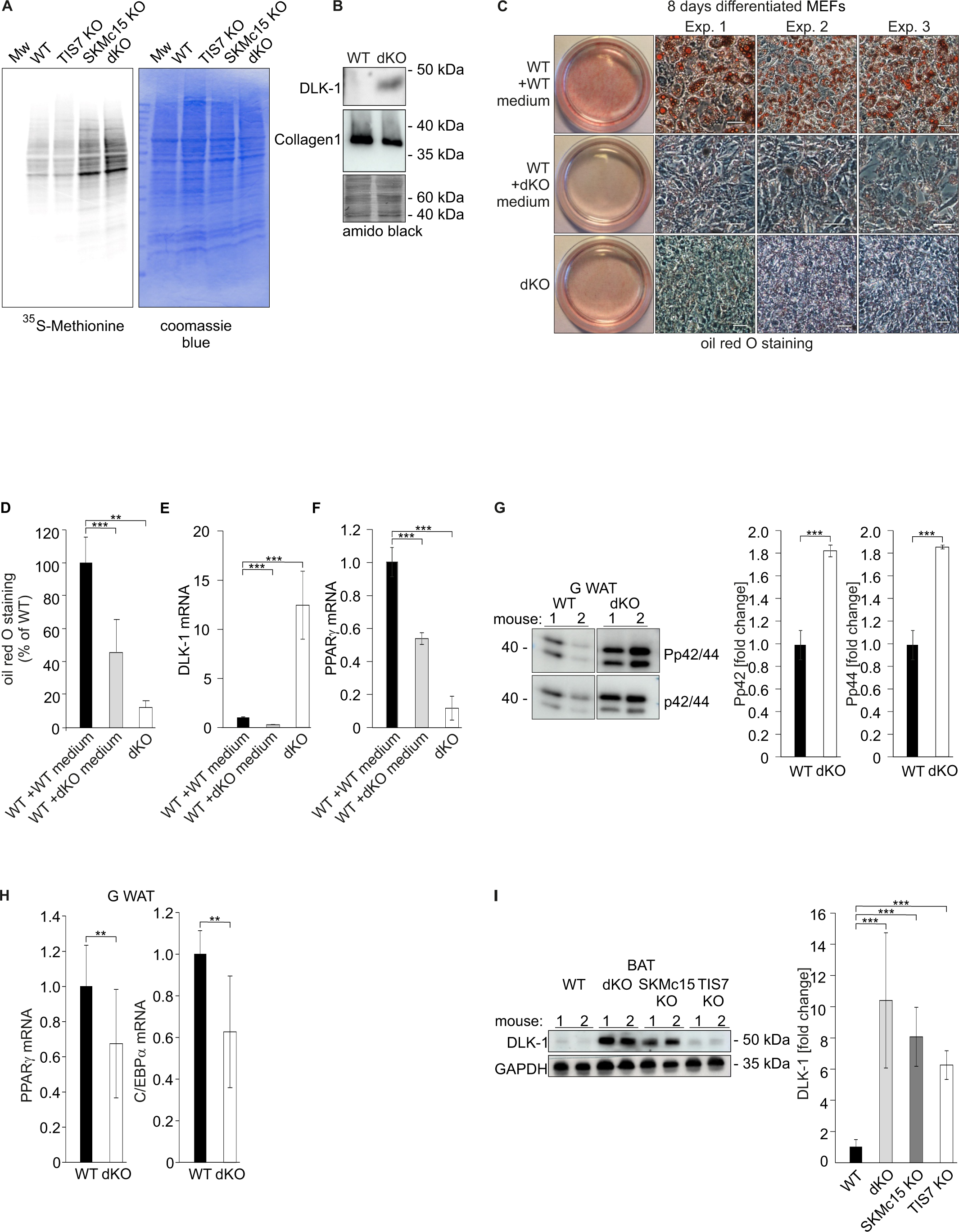
SKMc15 inhibits DLK-1 translation. dKO-secreted DLK-1 inhibits adipocyte differentiation through MEK/ERK signaling. A) Translational analysis – *in vivo* metabolic staining of MEFs with ^35^S-methionine. Equal numbers of cells were seeded and 24 hours later treated as explained in Methods section. Identical volumes of cell lysates were separated by SDS-PAGE, gel was dried and analyzed by a phosphorimager. Equal loading was documented by the Coomassie blue staining of the gel. B) dKO MEFs secrete DLK-1 protein into the cell culture medium. Identical volumes of media from WT and dKO MEF cells 8 days treated with the adipocyte differentiation cocktail were analyzed by Western blot. Collagen I was present in both samples in similar amounts. Equal loading of samples was evaluated by amido black staining of the membrane. C) dKO MEFs-conditioned medium inhibited adipocyte differentiation in WT MEFs. MEFs were treated 8 days with the adipocyte differentiation cocktail. WT MEFs treated with the dKO-conditioned medium (replaced 3 times every 2 days) showed reduced differentiation. Images representing three biological repeats were cropped and space bars represent always 20µm. D) Quantification of oil red O staining from three independent adipocyte differentiation experiments. Data shown are mean ± STD, ordinary one-way ANOVA P=0.0016. Student’s t-test, **P<0.01, *** P<0.001. E) DLK1 mRNA levels in WT and dKO MEFs treated 8 days with the adipocyte differentiation cocktail or in WT MEFs treated with the dKO-conditioned medium, *** P<0.001. F) PPARγ mRNA expression levels in same cells as shown in panel D and E, *** P<0.001. G) Representative Western blots of phospho-p44, phospho-p42, p44 and p42 in gonadal WAT of WT and dKO mice. Normalization on p44 and p42; n=3. Error bars indicate standard deviations. WT values were set as 1, *** P<0.001. H) PPARγ and C/EBPα mRNA expression in gonadal WAT. Normalization on GAPDH; n=3. Error bars indicate standard deviations. WT values were set as 1, ** P<0.01. I) DLK-1 protein is up regulated in brown adipose tissues (BAT) of SKMc15, TIS7 single knockout and in dKO mice. Western blot analysis was performed on 5 samples of each genotype. Normalization on GAPDH. Error bars indicate standard deviations, ***P<0.001.

### TIS7 and SKMc15 regulate adipocyte differentiation through DLK-1, MEK/ERK pathway, PPARγ and C/EBPα

DLK-1 protein carries a protease cleavage site in its extracellular domain (Lee, Helman et al., 1995) and is secreted. The extracellular domain of DLK-1 is cleaved by ADAM17, TNF-α Converting Enzyme to generate the biologically active soluble DLK-1 (Wang & Sul, 2009). DLK-1 mRNA and protein levels are high in preadipocytes but DLK-1 expression is absent in mature adipocytes. Hence, adding soluble DLK-1 to the medium inhibits adipogenesis (Garces, Ruiz-Hidalgo et al., 1999). To test if the dKO MEFs secreted DLK-1, cell culture media from MEFs treated 8 days with the adipocyte differentiation cocktail were collected and analyzed by Western blotting. As shown in Fig 4B, dKO cells secreted DLK-1 protein, but in an identical volume of the cell culture medium from WT MEFs no DLK-1 could be detected. In contrary to that, an unrelated secreted protein, namely Collagen I, was found in media of both, WT and dKO MEFs in similar, in the medium of WT MEFs even slightly higher amounts. To prove that secreted DLK-1 could inhibit adipocyte differentiation of dKO MEFs, we cultured WT MEFs with conditioned medium from dKO MEFs. Adipocyte differentiation of WT MEFs was strongly inhibited by the dKO MEFs-conditioned medium when compared to the control WT cells (Fig 4C, quantified in Fig 4D). Another indication that dKO inhibited adipocyte differentiation via DLK-1 protein up regulation delivered the experiment where we specifically knocked down DLK-1. We targeted either all Dlk1 mRNA splice variants (oligo shDLK-1 391) or only Dlk1 mRNA splice variants containing coding sequences for the protease site for extracellular cleavage (oligo shDLK-1 393) as previously published in (Mortensen et al., 2012). Both DLK-1 knockdown constructs stably expressed in dKO MEFs significantly increased (P<0.001) adipocyte differentiation as documented in Fig EV2C. Hes1 levels, together with DLK-1 are continuously down regulated during the process of adipogenesis while PPARγ are rising (Huang, Yang et al., 2010). In our experimental setup, down regulation of DLK-1 levels was paralleled by significant (P<0.001) decrease in Hes1 mRNA levels (Fig EV3A). Opposite, treatment with a recombinant DLK-1 protein or stable DLK-1 ectopic expression documented by qRT-PCR (Fig EV3D) significantly (P<0.001) inhibited adipocyte differentiation of WT MEFs as shown in Fig EV3B,C. This was accompanied by a significant (P<0.001) decrease in C/EBPα mRNA levels (Fig EV3E). Furthermore, DLK-1 mRNA quantification (Fig 4E) documented that dKO MEF cell lysates contained significant amounts of DLK-1 mRNA when compared to WT control or WT cells treated with dKO MEFs-conditioned medium. In parallel, WT cells treated with dKO MEFs-conditioned medium expressed significantly lower amounts of PPARγ mRNA when compared to WT cells incubated with control medium (Fig 4F). In addition, ectopic expression of SKMc15 and co-expression with TIS7 in dKO MEFs rescued almost up to WT levels the adipocyte differentiation potential of these cells (Fig EV2A). Ectopic expression of both, TIS7 and SKMc15 significantly (P<0.001) down regulated DLK-1 mRNA expression in dKO MEFs (Fig EV2A). Moreover, conditioned medium from dKO MEFs expressing DLK-1 shRNA knockdown constructs significantly (P<0.001) lost the ability to inhibit adipocyte differentiation of WT MEFs as the medium from dKO MEFs did (Fig EV3F). Based on these results we concluded that cells derived from dKO mice express increased DLK-1 levels and secreted, soluble DLK-1 may inhibit adipocyte differentiation *in vivo*.

Previous studies showed that soluble DLK-1 activates MEK/ERK signaling, which is required for inhibition of adipogenesis (Kim et al., 2007). As DLK-1 was strongly up regulated in dKO SVF cells during adipocyte differentiation we subsequently analyzed possible activation of the MEK/ERK pathway in gonadal WAT samples of WT and dKO mice (Fig 4G). The phosphorylation of p44 and of p42 was up regulated 1.8-fold (P<0.001) in the dKO when compared to the WT G WAT samples (Fig 4G). Next, we addressed the question of expression levels of PPARγ and C/EBPα in WAT depots. Both adipocyte differentiation regulators PPARγ and C/EBPα mRNA expression levels were in gonadal WAT samples isolated from dKO mice significantly (P<0.01) downregulated when compared to the values of WT control animals (Fig 4H). Furthermore, we analyzed the possible effect of TIS7, SKMc15 and their combined knockout on the DLK-1 levels in brown adipose tissue (BAT). Western blot analysis identified significant (P<0.001) increase in DLK-1 protein levels in BAT samples in knockout mice of all three genotypes (Fig 4I). The MEK/ERK pathway was similarly as in gonadal WAT, up regulated also in MEFs generated from dKO mice (Fig 5A). The phosphorylation of p42 and of p44 was up regulated 3-fold and 4.3-fold, respectively (P<0.01), in the adipocyte-differentiated dKO when compared to the WT MEFs (Fig 5B). Previously, activation of MEK/ERK by DLK-1 was shown to up regulate the expression of the transcription factor SOX9, resulting in the inhibition of adipogenesis (Sul, 2009). Therefore, we measured SOX9 mRNA expression by RT qPCR in 8 days adipocyte-differentiated MEFs. SOX9 expression was significantly (P<0.001) up regulated in 8 days adipocyte-differentiated dKO when compared to WT MEFs (Fig 5C). PPARγ and C/EBPα mRNA levels were both continuously up regulated during the 8 days differentiation protocol of WT MEFs (Fig 5D,E). On the contrary, we did not find any significant increase in PPARγ and C/EBPα mRNA levels in dKO MEFs (Fig 5D,E). In a rescue experiment we showed that only the co-expression of TIS7 and SKMc15 strongly increased the expression of the PPARγ in undifferentiated dKO MEFs (P<0.001), almost up to the levels of WT MEFs (Fig 5F). The expression of C/EBPα was also strongly up regulated by the co-expression of TIS7 and SKMc15 (P<0.001) (Fig 5G). Based on these results we concluded that TIS7 and SKMc15 regulate the expression of both PPARγ and C/EBPα, crucial regulators of adipocyte differentiation.

**Figure 5.**
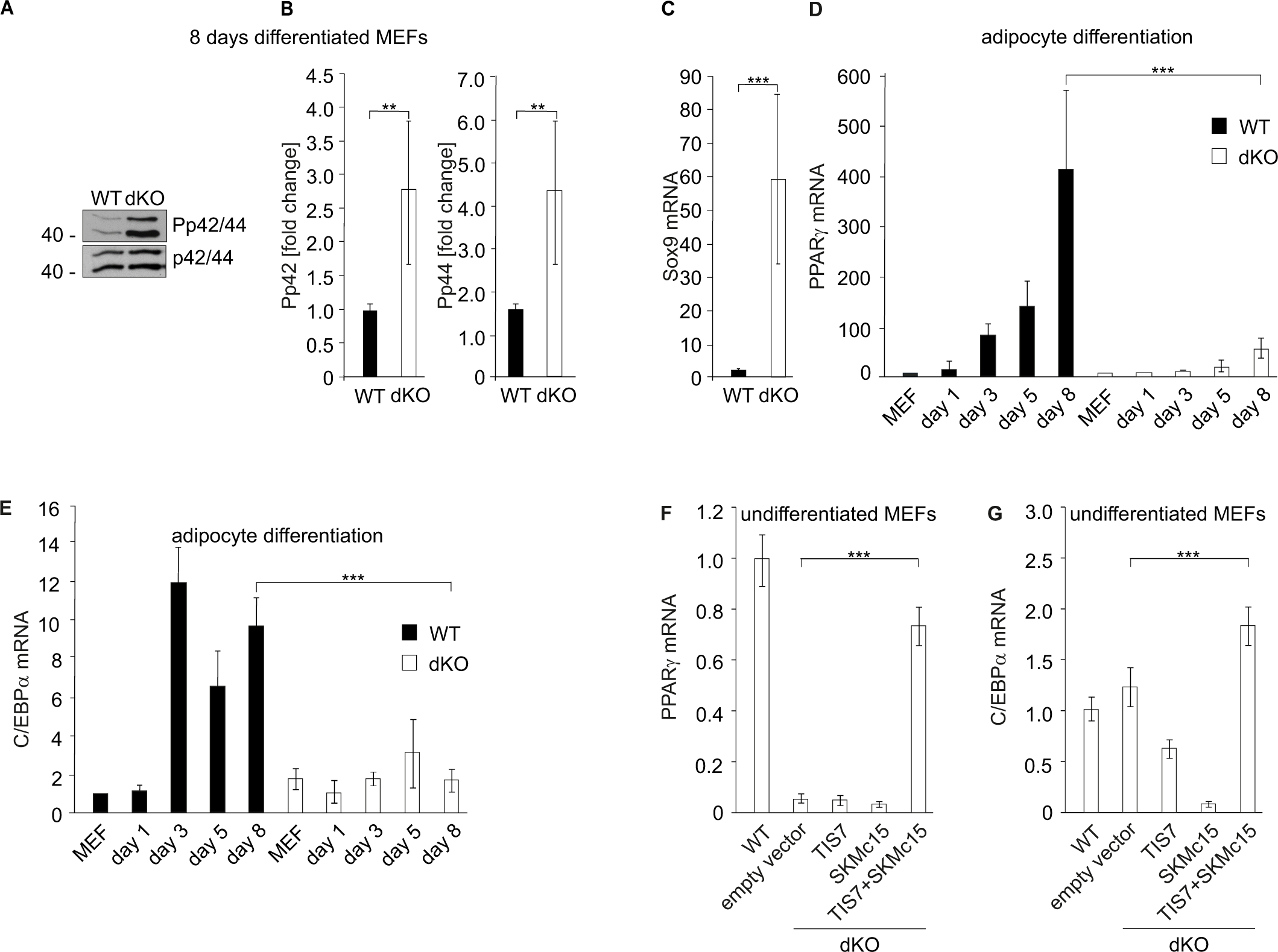
MAPK signaling and Sox9 expression are induced while adipocyte differentiation regulatory genes are down regulated in dKO MEFs. A) Representative Western blots of phospho-p44, phospho-p42, p44 and p42 from 8 days adipocyte differentiation cocktail-treated MEFs. B) Quantitative analysis of Western blot data. Normalization on p44 and p42; n=3. Error bars indicate standard deviations. WT values were set as 1; **P<0.01. C) SOX9 expression was measured by qPCR in 8 days adipocyte-differentiated WT and dKO MEFs. SOX9 values were normalized on GADPH expression. WT MEFs values were set as 1, n=3. Data shown are mean ± STD. Student’s t-test ***P<0.001. D) PPARγ and E) C/EBPα mRNA levels were down regulated in dKO adipocyte-differentiated MEFs. Gene expression was measured by qPCR during the treatment with the adipocyte differentiation cocktail. Values were normalized on GADPH expression. WT MEFs values were set as 1, n=3. Data shown are mean ± STD, *** P<0.001. F) Ectopic co-expression of TIS7 and SKMc15 in undifferentiated MEFs significantly increased levels of adipogenic genes PPARγ and G) C/EBPα. Data shown are mean ± STD, Student’s t-test ***P<0.001.

### Lipid absorption is reduced in TIS7 and SKMc15 double knockout mice

dKO mice were despite identical food intake, activity and RER (Suppl fig 1A and Fig EV1B) leaner than their WT littermates. Therefore, we tested the possibility that dKO mice store energy ectopically. In the feces of dKO mice fed with high fat diet we identified significantly higher (P<0.05) amounts of free fatty acids than in that of their WT siblings (Fig 6A). Secondly, the energy content of dried feces from dKO mice determined by bomb calorimetry was significantly higher (P<0.001) than that of WT mice (Fig 6B). PPARγ induces the expression of CD36, a very long chain fatty acids (VLCFA) transporter in heart, skeletal muscle, and adipose tissues (Coburn, Knapp et al., 2000). The regulation of CD36 by PPARγ contributes to the control of blood lipids. Interestingly, CD36 null mice exhibit elevated circulating LCFA and triglycerides levels consistent with the phenotype of dKO mice and CD36 deficiency partially protected from HFD-induced insulin resistance (Wilson, Tran et al., 2016). Because of down regulated PPARγ levels in adipose tissues of dKO mice we decided to study more in detail CD36 regulation of these as well. In inguinal WAT from dKO mice, we found strongly reduced CD36 mRNA expression levels (64 % of the WT values) (Fig 6C). Next, we analyzed CD36 in WT and dKO MEFs before onset and during adipocyte differentiation. As long as CD36 mRNA expression substantially increased in WT MEFs, there were almost undetectable transcript levels of CD36 in dKO cells treated with the adipocyte differentiation cocktail (Fig 6D). Diacylglycerol acyltransferase 1 (DGAT1), a protein associated with the enterocytic triglyceride absorption and intracellular lipid processing (Nozaki, Tanaka et al., 1999) is besides CD36 another target gene of adipogenesis master regulator PPARγ (Koliwad, Streeper et al., 2010). DGAT1 mRNA levels are strongly up regulated during adipocyte differentiation (Cases, Smith et al., 1998), its promoter region contains a PPARγ binding site (Ludwig, Mahley et al., 2002) and DGAT1 is also negatively regulated by the MEK/ERK pathway (Tsai, Qiu et al., 2007). DGAT1 expression was shown to be increased in TIS7 transgenic mice (Wang et al., 2005) and its expression was decreased in the gut of high fat diet-fed TIS7 KO mice (Yu et al., 2010). Importantly, DGAT1 expression in adipocytes and inguinal WAT is up regulated by PPARγ activation (Koliwad et al., 2010). Therefore, we measured DGAT1mRNA levels during the differentiation of WT and dKO MEFs into adipocytes to analyze the role of TIS7 and SKMc15 on the regulation of this protein involved in adipogenesis and triglyceride processing. As long as DGAT1 expression substantially increased during the differentiation of WT MEFs, there was no difference in DGAT1 mRNA levels in dKO cells treated with the adipocyte differentiation cocktail (Fig 6E). We concluded that TIS7 and SKMc15 regulate expression of multiple proteins involved both, in adipocyte differentiation and in fat uptake, thereby contributing to the lean phenotype of dKO mice through multiple means.

**Figure 6.**
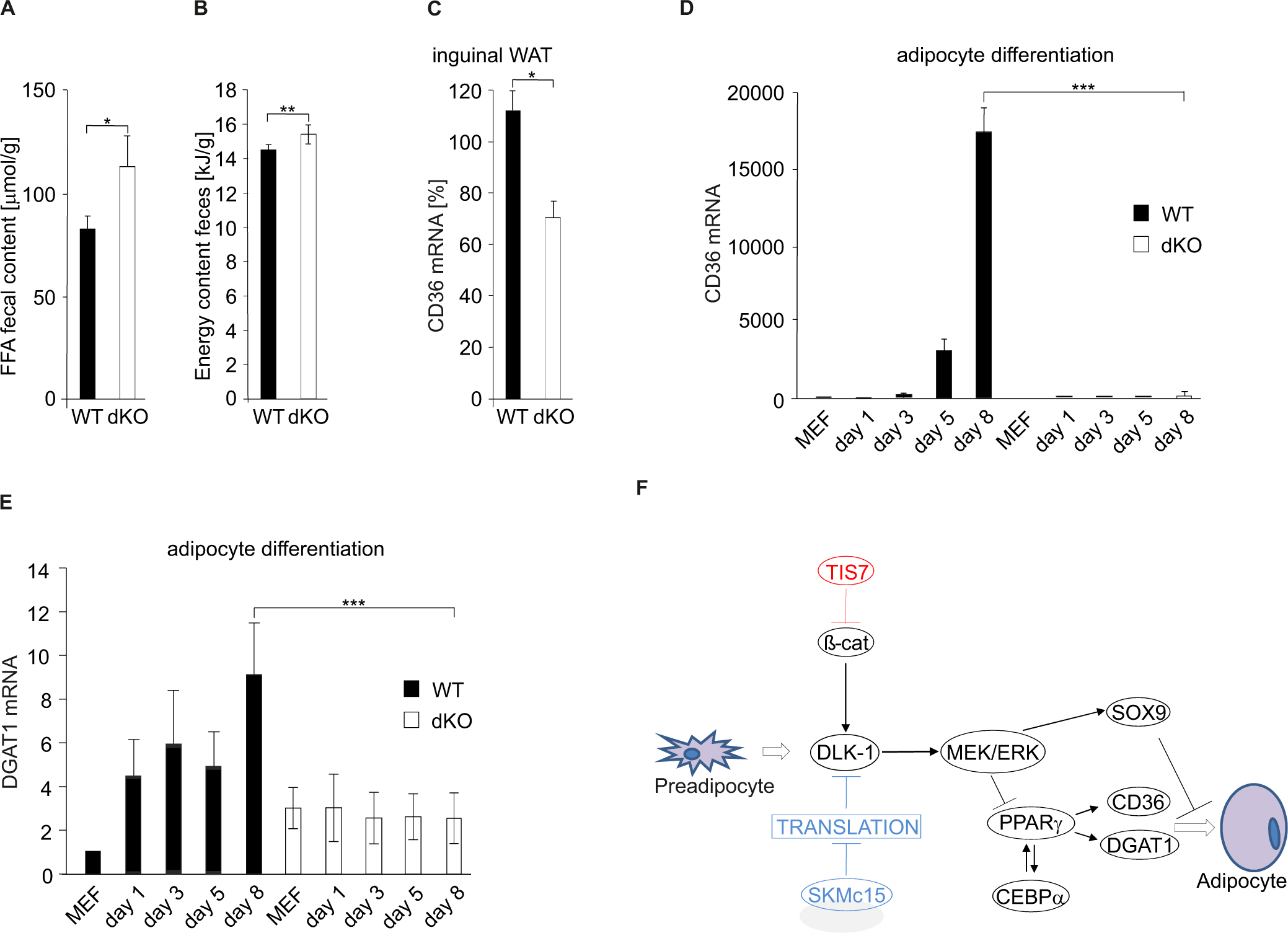
FFA uptake is inhibited in dKO mice. A) Free fatty acids concentrations in feces of WT and dKO mice. Feces were collected every second day and the composition of excreted lipids was determined by capillary gas chromatography. Data shown are mean ± STD, *P<0.05. B) Energy content of dried egested feces samples was determined by bomb calorimetry. Data shown are mean ± STD, **P<0.01. C) qPCR analysis of CD36 mRNA levels in inguinal WAT. Data shown are mean ± STD, *P<0.05. D) qPCR analysis of CD36 in MEFs differentiating into adipocytes. Relative expression levels were normalized on GAPDH expression. WT MEFs values were set as 1. Data shown are mean ± STD, ***P<0.001. E) DGAT1 mRNA levels were down regulated in dKO adipocyte-differentiated MEFs. Data shown are mean ± STD, ***P<0.001. F) Proposed model of TIS7 and SKMc15 molecular mechanisms of action during adipocyte differentiation. Two parallel mechanisms leading to a deficiency in adipocyte differentiation: TIS7-regulated Wnt signaling affected Dlk-1 transcription and SKMc15 acting as a translational inhibitor also contributed to the regulation of adipogenesis.

## Discussion

In this study, we show that simultaneous depletion of TIS7 and of its orthologue SKMc15, a protein recently identified as a translational inhibitor, caused severe reduction of adipose tissues and resistance against high fat-induced obesity in mice. We identified two parallel mechanisms leading to a deficiency in adipocyte differentiation. Firstly, TIS7-regulated Wnt signaling affected Dlk-1 transcription and secondly, our study provided the first experimental evidence for SKMc15 acting as a translational inhibitor that among other proteins controls DLK-1 protein levels, thereby contributing to the regulation of adipogenesis (Fig 6F).

dKO mice were phenotypically similar with both, CD36 deficient and DLK-1 transgenic mice, namely in decreased amounts of WAT and resistance to HFD-induced obesity. Previous studies showed that overexpression of TIS7 caused increased intestinal lipid transport resulting in elevated body weight gain during HFD feeding (Wang et al., 2015). Knockout of TIS7 and SKMc15 impaired absorption of free fatty acids from the lumen into enterocytes and reduced rate of fat absorption from intestines into the circulation. Simultaneously, higher concentrations of free fatty acids in the feces of dKO mice (Fig 6A) suggested that mice lacking TIS7 and SKMc15 suffer from an intestinal lipid uptake deficiency, yet another contributing reason for the lean phenotype of these mice. In dKO mice, we found inhibited PPARγ and C/EBPα levels, induced MEK/ERK pathway, and decreased expression of CD36 and DGAT1, all hallmarks of up regulated DLK-1, a known adipogenesis inhibitor. TIS7 and SKMc15 acted after the commitment to the preadipocyte stage since the elevated DLK-1 and β-catenin levels found in WAT of dKO mice were characteristic for the preadipocyte stage (Gautam, Khedgikar et al., 2017).

It is known that disruption of Wnt signaling in embryonic fibroblasts results in spontaneous adipocyte differentiation (Bennett, Hodge et al., 2003) and opposite, stabilized β-catenin keeps cells in the preadipocyte stage (Ross et al., 2000). A subpopulation of the stromal vascular fraction of adipose tissue is adipogenic and at the same time has a weaker Wnt/β-catenin signal (Hu, Yang et al., 2015). Consistent with this knowledge we found up regulated β-catenin protein levels and increased Wnt signaling activity in WAT of TIS7 single and dKO MEFs. It has to be mentioned that our data were different to those by Nakamura *et al*. who showed that following the TIS7 overexpression in adipocytes was up regulated Wnt/β-catenin signaling and inhibited oil red O staining (Nakamura et al., 2013). However, one has to take into consideration the difference in cell systems used in these two studies. All results presented in Nakamura’s study were obtained in 3T3-L1 cells fibroblasts following overexpression of TIS7. Opposite to that, we have studied the role of endogenous TIS7 in cells derived from wild type or knockout mice. Moreover, we have found ubiquitous TIS7 expression in all WT mouse organs without any pre-treatment such as e.g. hypoxia. On the other hand, Nakamura *et al*. identified up regulated TIS7 expression levels in WAT of obesity model mice. This result, however, fully supports our findings of lean phenotype in TIS7 and SKMc15 dKO mice.

We demonstrated that in dKO MEFs, induced for adipocyte differentiation, the increase in Wnt signaling led to up regulated binding of β-catenin to the DLK-1 gene regulatory elements, resulting in sustained DLK-1 expression. In adipogenesis DLK-1 expression is down regulated through histone methylation by COPR5 and PRMT5 that prevent β-catenin binding to the DLK-1 gene (Paul et al., 2015). Interestingly, in dKO MEFs we found weaker binding of dimethylated H4R3 to the DLK-1 gene that was consistent with our previous findings on the negative role of TIS7 in epigenetic regulation of gene expression including the PRMT5 activity (Lammirato et al., 2016).

There was a difference in effects of ectopic expression of TIS7, SKMc15 and their co-expression on DLK-1 levels (Fig 3B), confirming the hypothesis that TIS7 and SKMc15 regulate DLK-1 levels via two independent pathways/mechanisms. Despite the fact that SKMc15 knockout had no effect on Wnt signaling it nevertheless affected DLK-1 levels, suggesting a contribution of SKMc15 to its regulation. Since SKMc15 was identified as a novel factor translationally inactivating ribosomes (Brown et al., 2018) and given that down regulation of the protein synthesis machinery is an essential regulatory event during early adipogenic differentiation (Marcon et al., 2017), we proposed that SKMc15 regulates DLK-1 levels through translational regulation. Experiments presented here confirmed specific regulation of DLK-1 through this mechanism. Elevated DLK-1 levels in dKO MEFs activated the MEK/ERK pathway thereby decreasing PPARγ and C/EBPα levels important for adipogenic differentiation. Besides that, the expression of Sox9 and Hes1 were up regulated, suggesting that the dKO MEFs keep their proliferative state and cannot enter differentiation into mature adipocytes (Kim et al., 2007).

Down regulation of PPARγ and C/EBPα implied changes in expression of downstream adipogenesis-related genes. Among them, decreased CD36 and DGAT1 levels were a plausible explanation of the dKO mice lean phenotype since it is known that both proteins play a functional role in differentiation of murine adipocytes and their deficiency impairs fat pad formation independent of lipid uptake (Christiaens, Van Hul et al., 2012). Our findings implicate that TIS7 and SKMc15 play a up to now unknown role in the regulation of CD36 and therefore possibly contribute to CD36 related pathogenesis of human metabolic diseases, such as hypoglycemia (Nagasaka, Yorifuji et al., 2011), hypertriglyceridemia (Kashiwagi, Tomiyama et al., 2001), and disordered fatty acid metabolism (Glazier, Scott et al., 2002, Tanaka, Nakata et al., 2001).

We and others have previously shown that TIS7 is involved under both, chow and HFD conditions in intestinal triglyceride absorption. Here we show for the first time the role of TIS7 and its orthologue SKMc15 in adipocyte differentiation. Moreover, our data presented here confirm the physiological function of SKMc15 as a translational inhibitor. To our surprise, although TIS7 and SKMc15 share high sequence homology and a functional role in the adipogenesis, they use two independent regulatory mechanisms.

## Methods

### Generation of animal models

All animal experiments were performed in accordance with Austrian legislation BGB1 Nr. 501/1988 i.d.F. 162/2005).

Mice lacking TIS7 were previously described (Vadivelu et al., 2004). SKMc15 KO mice generation: targeting construct contained SKMc15 gene locus exons. A loxP site and a neomycin resistance gene inserted at position 105515 (AY162905). The neomycin cassette was flanked by two frt sites. A second loxP site was inserted at position 102082 (AY162905). Further downstream, 15 additional nucleotides, part of intron 1, exon 2 (splice site: donor and acceptor) and the CDS of hrGFP from the Vitality hrGFP mammalian expression vector pIRES-hrGFP-2a (Stratagene) were added. The targeted construct was electroporated into Sv129 mouse ES cells. After the selection with G418 single cell clones were screened by PCR and confirmed by Southern blot analysis. After Cre recombinase treatment cell clones were screened by PCR. Clones with floxed gene deleted, were used for blastocyst injection into C57Bl/6J mice. Two male chimeras with ≥ 40 % or more agouti coat color were mated to C57Bl6 females. The knockout mouse strain was derived from one male mouse carrying the allele of interest. Heterozygous mice were back-crossed to C57Bl6 mice for 9 generations. dKO mice were generated by crossing the SKMc15 KO mice with the TIS7 single KO mice. The resulting mice were screened by PCR and double heterozygous mice used for further breeding until homozygosity. In order to achieve maximal homogeneity of experimental groups, in all experiments presented here we used only male mice.

### SKMc15 knockout Southern blot analysis

XbaI restriction sites (position 109042; position 102079; position 92253) located in the SKMc15 locus (AY162905). Fragment detected by the Southern probe: 9.6 kb wt (Suppl fig 1F, insert, band V); 15 kb in deleted locus (band IV). The 566 bp long probe for hybridization from genomic DNA (96605 – 97171; AY162905). PCR primer sequences: RK 150 Fwd: 5’-GGTCCTGCCACTAATGCACTG-3’; RK 151 Rev: 5’-GCAGACAGATGCCAGGAAGAC-3’.

### SKMc15 knockout PCR genotyping analysis

hrGFP insert in the 3’ UTR detected by primer GFP2 5’-AGCCATACCACATTTGTAGAG-3’ and RK101 3’ UTR (5’-TGATGATAGCTTCAAAGAGAA-3’; 100617 – 100591 of the SKMc15 locus (AY162905). PCR product 1700 bp. SKMc15 detection: RG1; 5’-TGTGGCCTTTATCCTGAGTC-3’; 102286–102266) and RG2; 5’-TGGCTTCATTTACACTACTCCTT-3’; 101860 – 101882 primers (Suppl fig 1F). WT allele PCR product 426 bp and the targeted allele PCR product 1772 bp (Suppl fig 1G). TIS7 genotype tested as explained previously (Vadivelu et al., 2004) (Suppl fig 1H).

### Growth monitoring and body composition measurement

Mice weaned at 3 weeks, regular chow diet, weighed weekly. Dual energy X-ray absorptiometry was measured with Norland scanner (Fisher Biomedical). Micro-CT experiments performed using vivaCT 40 (Scanco Medical AG). The scans were performed using 250 projections with 1024 samples, resulting in a 38µm isotropic resolution. Tube settings: 45kV voltage, 177µA current, integration time 300ms per projection. Image matrix 1024*1024 voxels and a grayscale depth of 16 bit. The length of the image stack was individually dependent, starting from the cranial end of the fist lumbar vertebrae, to the caudal end of the fifth lumbar vertebrae. The image reconstruction and post processing were performed using the Scanco Medical system software V6.6. For the adipose tissue evaluation an IPL (image processing language) script by Judex et al., provided by Scanco medical AG was modified to the scanner individual parameters leading to 2 values lower threshold than in the original script for the adipose tissue filters 76. The script calculated the total abdominal volume without potential air in the cavities. A separation of subcutaneous and visceral fat mass was used only for visualization. For the quantitative fat mass analysis we used 56 male 5 to 12 month old mice (mean age 9.35 ± 2.03 month) reflecting an adult to mid aged cohort defined by Flurkey et al. 2007 (Neeland, Ayers et al., 2013). The development of adipose tissue in healthy mice is stable between 4 and 12 months (Lemonnier, 1972). Therefore, sex and age could have only very limited influence on the experimental results. For the quantitative comparison, the percent contribution of the abdominal fat to the body weight was calculated using a mean weight of 0.9196 g/ml for adipose tissue (Neeland et al., 2013). Statistics were performed using an ANCOVA with a Bonferroni corrected Post Hoc testing on the µCT fat data and the body mass data.

### Metabolic measurements

The indirect calorimetry trial monitoring gas exchange, activity, and food intake was conducted over 21 hours (PhenoMaster TSE Systems). Body mass and rectal body temperature before and after the trial were measured. The genotype effects were statistically analyzed using 1-way ANOVA. Food intake and energy expenditure were analyzed using a linear model including body mass as a co-variate.

### HFD feeding

Age-matched (7-week-old) male TIS7-/-SKMc15-/-and WT mice were caged individually and maintained up to 3 weeks on a synthetic, high saturated fat (HFD) diet (TD.88137; Ssniff). Animals were weighted every 4th day between 08:00 and 10:00. Small intestines were harvested for oil red O staining to detect lipid accumulation. Intestines, liver, muscles and adipose tissue were collected for total RNA and protein isolation. Unfixed intestines were flushed with PBS using a syringe, embedded in Tissue-Tek (Sakura, 4583) and frozen in liquid nitrogen for immunohistochemical analysis.

### Quantitative food consumption and fecal fat determination

Adult mice were acclimatized to individual caging and to the HFD for a week, monitored for weight gain and their food intake daily. The daily food intake data were pooled for the following 7 days and the food intake was estimated (g/day). Feces were collected daily and weighted for 7 days after second week of the HFD consumption. 50 mg of dried feces were boiled in 1 ml alkaline methanol (1M NaOH/Methanol, 1 3 v/v) at 80°C for 2 h after addition of 50 nmol 5α-Cholestane (Sigma, C8003) as internal standard for neutral sterol analysis. After cooling down to room temperature, neutral sterols were extracted using three times 3 ml of petroleum ether, boiling range 60–80°C. The residual sample was diluted 1:9 with distilled water. 100 µl of the solution were subjected to an enzymatic total bile acid measurement. The extracted neutral sterols were converted to trimethylsilyl derivatives. Neutral sterol composition of prepared feces samples was determined by capillary gas chromatography on an Agilent gas chromatograph (HP 6890) equipped with a 25 m×0.25 mm CP-Sil-19 fused silica column (Varian) and a Flame Ionization Detector. The working conditions were the following: Injector temperature 280°C; pressure 16.0 psi; column flow constant at 0.8 ml/min; oven temperature program: 240°C (4 min), 10°C/min to 280°C (27 min); detector temperature 300°C (Harchaoui, Franssen et al., 2009). Energy content of dried egested feces samples (∼ 1 g per sample) was determined by bomb calorimetry (IKA C 7000, IKA, Staufen, Germany) (Pfluger, Kabra et al., 2015).

### Antibodies, viral and cDNA constructs

Antibodies: anti-CD36 AbD Serotec (MCA2748), Abcam (ab36977), p44/42 MAPK (Erk1/2) and Phospho-p44/42 MAPK (Erk1/2) (Thr202/Tyr204) Cell Signaling Technology (9102, 9101), β-catenin antibody Sigma (C2206), anti-DLK Abcam (ab119930), anti-histone H4R3me2s antibody Active Motif (61187). For ChIP experiments were used anti-TIS7 (Vietor et al., 2002) and anti-SKMc15 (Vadivelu et al., 2004) rabbit polyclonal antibodies previously proven for ChIP suitability in (Lammirato et al., 2016). pTOPflash reporter construct was a gift from H. Clevers (University of Utrecht, Holland). TIS7 construct was described previously (Vietor et al., 2002). Partial cds of mSKMc15 was amplified by PCR and cloned into pcDNA3.1(-)/MycHis6 (Invitrogen).

### Cell culture and adipocyte differentiation

MEFs were generated from 16-day old embryos. After dissection of head for genotyping, removal of limbs, liver and visceral organs embryos were minced and incubated in 1 mg/mL collagenase (Sigma-Aldrich, C2674) 30 mins at 37°C. Embryonic fibroblasts were maintained in growth medium containing DMEM high glucose (4.5 g/l); sodium pyruvate, L-glutamine, 10% FCS (Invitrogen, 41966029), 10% penicillin/streptomycin at 37°C in 5% CO_2_. Adipocyte differentiation treatment was medium with 0.5 mM 3-isobutyl-1-methylxanthine (I5879), 1 µM dexamethasone (D4902), 5 µg/ml insulin (I2643) and 1 µM Rosiglitazone (Sigma-Aldrich, R2408). After 3 days growth medium containing 1 µg/ml insulin cells were differentiated for 8 days. To visualize lipid accumulation adipocytes were washed with PBS, fixed with 6 % formaldehyde overnight, incubated with 60 % isopropanol, air dried and then incubated with oil red O. Microscopic analysis was followed by isopropanol elution and absorbance measurement at 490 nm.

### Construction of expression plasmids and generation of stable cell lines

The pRRL CMV GFP Sin-18 plasmid (Zufferey et al., 1998) was used to generate TIS7, SKMC15 and Dlk1 expression lentiviral constructs. For this, the corresponding cDNAs were cloned into the BamHI and SalI sites of the pRRL CMV GFP Sin-18 plasmid. A cap-independent translation enhancer (CITE) fused to the puromycin resistance pac gene and the woodchuck hepatitis virus post-transcriptional regulatory element (WPRE) were introduced downstream of the TIS7, SKMC15 and Dlk1 coding sequences. All DNA constructs were verified by sequencing.

To generate the Dlk1 shRNA lentiviral vectors, oligonucleotides targeting either all Dlk1 mRNA splice variants (Dlk1Total: 5’-GATCCCCAGATCGTAGCCGCAACCAATTCAAGAGATTGGTTGCGGCTACGATCTTTTTTGGAAA-3‘) or only Dlk1 mRNA splice variants containing the extracellular cleavage sequence (Dlk1PS: 5’-GATCCCCTCCTGAAGGTGTCCATGAATTCAA GAGATTCATGGACACCTTCAGGATTTTTGGAAA-3‘) (Mortensen et al., 2012) were fused with the H1 promoter and cloned into the pRDI292 vector as reported (Reintjes et al., 2016). The GFP-targeting pRDI-shRNA-GFP plasmid was used as a control (Reintjes et al., 2016).

The viral supernatants were obtained as described previously (Leitner, Jakschitz et al., 2022), concentrated with Retro-X Concentrator (Clontech, Takara Bio) and used to infect WT, TIS7 and SKMc15 single and dKO MEFs. The selection was carried out for 2 weeks in DMEM supplemented with 10% (v/v) FBS, 100 U mL^−1^ penicillin and 100 μg mL^−1^ streptomycin, and 2 µg mL^−1^ puromycin.

### Polysome profiling

Polysome profiling was performed as described in (Savant-Bhonsale & Cleveland, 1992), with modifications. The day before harvesting the cells, continuous 15-45 % (w/v) sucrose gradients were prepared in SW41 tubes (Beckman) in Polysome Gradient Buffer (10 mM Hepes-KOH, pH 7.6; 100 mM KCl; 5 mM MgCl2) employing the Gradient Master ip (Biocomp) and stored overnight at 4 °C. All steps of the protein extraction were performed on ice. Exponentially growing cells were washed twice with ice cold DPBS (Gibco) supplemented with 100 µg/mL f. c. cycloheximide, scraped in 300 µL Polysome Lysis Buffer (10 mM Hepes-KOH, pH 7.6; 100 mM KCl; 5 mM MgCl2; 0.5 % IGEPAL® CA-630; 100 µg/mL cycloheximide) supplemented with 0.1 U/µL murine RNase Inhibitor (NEB), and passed through a G25 needle 25 times. Nuclei were pelleted at 16,000 g for 6 min at 4 °C and the supernatants were carefully layered onto the sucrose gradients. Samples were centrifuged at 35,000 rpm for 2 h at 4 °C (with brakes switched off) using an SW 41 Ti rotor (Beckmann). 20 fractions of 0.6 mL were collected by use of a peristaltic Pump P1 (Amersham Biosciences) and polysome profiles were generated by optical density measurement at 254 nm using optical unit UV-1 (Amershan Biosciences) and chart recorder Rec 111 (Amersham Biosciences).

### Analysis of translation by metabolic labelling

Pulse labelling of mitochondrial proteins was performed as described before (Popow, Alleaume et al., 2015), with the following changes: 1x 10^6^ wild-type and dKO MEFs cells per plate were seeded and cultivated for 24 h. Cells were washed twice with PBS followed by one wash with labelling medium (ESC medium without cysteine and methionine) and a 30 min incubation in labelling medium. To label newly translated products, 200 µCi of [^35^S]-methionine (10 mCi/ml; Hartman Analytic) were added and the cells were incubated for 1 h. Cells were washed and incubated another 10 min in standard medium at 37 °C. Cells were then harvested by trypsinizing, washed once with ice-cold PBS, extracted with ice-cold RIPA buffer containing protease-inhibitors and briefly sonicated. Proteins were fractionated by gel electrophoresis in 16% Tricine gels (Invitrogen) and stained with Coomassie brilliant blue, and radioactive signals were visualized by phosphorimaging. Signal intensities were quantified using the Image Studio Lite (v5.2) software.

### RT-PCR

Tissues from animals fed for 3 weeks with HFD were snap frozen and stored at –80°C. Total RNA was isolated using the TRIzol reagent (Invitrogen, 15596026). RNA was then chloroform-extracted and precipitated with isopropanol. The yield and purity of RNA was determined by spectroscopic analysis; RNA was stored at –80°C until use.

### Quantitative RT-PCR and statistics

Total RNA were treated with DNAse1 and reverse transcribed to cDNA by Revert Aid First Strand cDNA Synthesis Kit (Thermo Scientific, K1622) with oligo dT primers. Quantitative RT-PCR was performed using Taqman probes and primer sets (Applied Biosystems) specific for CD36 (assay ID Mm00432398_m1), Dgat1 (Mm00515643_m1), Pparγ (Mm00440940_m1), C/EBPα (Mm00514283_s1), DLK-1 (Mm00494477_m1) and Sox9 (Mm00448840_m1). Ribosomal protein 20 (assay ID Mm02342828_g1) was used as normalization control for quantification by the ddCt method. PCR reactions were performed using 10 µl cDNA in PikoReal 96 real-time PCR system (Thermo Scientific). Quantification data were analyzed by 2-tailed, Homoscedastic t-tests are based on the assumption that variances between two sample data ranges are equal type 2 Student’s t test.

### Transient transfections and luciferase assay

pGL2-Basic (Promega, E1641) or pGLCD36 (Shore, Dietrich et al., 2002) were used as reporter constructs. Expression constructs or empty vector DNA as a control were co-transfected. pCMV-β-Gal plasmid was used to normalize for transfection efficiency. For luciferase reporter assays, 1.5×105 cells were seeded into 24-well plates and transfected after 24 hours with indicated plasmid combinations using Lipofectamine Plus™ Reagent (Invitrogen, 15338030). The total amount of transfected DNA (2 μg DNA per well) was equalized by addition of empty vector DNA. Cells were harvested 48 h post-transfection in 0.25 M Tris, pH 7.5, 1% Triton X-100 buffer and assayed for both luciferase and β-galactosidase activities. Luciferase activity and β-galactosidase activity were assayed in parallel by using the Lucy 2® detection system (Anthos). Transfections were performed in triplicates and all experiments were repeated several times.

### Chromatin immunoprecipitation (ChIP)

Chromatin was isolated from TIS7 WT and dKO formaldehyde-treated, 8 days adipocyte differentiated MEFs using the EpiSeeker Chromatin Extraction Kit (Abcam, ab117152). ChIP analyses were carried out as described previously (Reintjes et al., 2016). Sequence of the oligonucleotides for two regions of the Dlk-1 promoter, encompassing TCF and β-catenin binding sites were as defined in (Paul et al., 2015). Sonicated chromatin was centrifuged at 15.000x g for 10 min at 4°C, and the supernatant (65 µg of sheared DNA per each IP) was diluted ten-fold with cold ChIP dilution buffer containing 16.7 mM Tris-HCl pH 8.1, 167 mM NaCl, 0.01% (w/v) SDS, 1.1% (w/v) Triton X-100 and 1.2 mM EDTA with protease inhibitors. Samples were pre-cleared for 1 h with protein A Sepharose CL-4B (Sigma-Aldrich, 17-0780-01) beads blocked with 0.2 µg/µl sonicated herring sperm DNA (Thermo Fisher, 15634017) and 0.5 µg/µl BSA (NEB, B9000 S). Immunoprecipitations were performed at 4°C overnight. Immune complexes were collected with protein A Sepharose for 1h at 4°C followed by centrifugation at 1000 rpm and 4°C for 5 min. Beads were washed with 1ml low salt wash buffer (20 mM Tris-HCl pH 8.1, 150 mM NaCl, 0.1%(w/v) SDS, 1%(w/v) Triton X-100 (Merck), 2 mM EDTA), high salt wash buffer (20 mM Tris-HCl pH 8.1, 500 mM NaCl, 0.1% (w/v) SDS, 1% (w/v) Triton X-100, 2 mM EDTA), LiCl wash buffer (10 mM Tris-HCl pH 8.1, 250 mM LiCl, 1% (w/v) sodium deoxycholate, 1% (w/v) IGEPAL-CA630, 1 mM EDTA) for 5 min at 4°C on a rotating wheel, and twice with 1 ml TE buffer (10 mM Tris-HCl pH 8.0, 1 mM EDTA). Protein-DNA complexes were eluted from antibodies by adding a freshly prepared elution buffer containing 1% SDS and 0.1 M NaHCO_3_. The eluate was reverse cross linked by adding NaCl to a final concentration of 0.2 M and incubating at 65 °C for four hours. Afterwards the eluate was treated with Proteinase K at 45 °C for 1 hour. The immunoprecipitated DNA was then isolated by phenol/chloroform precipitation and used as a template for real-time-quantitative PCR. The primer pairs specific for regulatory regions of the Dlk-1 gene were selected as described before (Paul et al., 2015). Reactions with rabbit IgG or with 1.23% of total chromatin (input) were used as controls. For real-time-quantitative PCR a PikoReal System was used. Signals were normalized to input chromatin and shown as % input. The raw cycle threshold (Ct) values of the input were adjusted to 100% by calculating raw Ct – log2(100/input). To calculate the % input of the immunoprecipitations, the equation 100 × 2[Ct (adjusted input to 100%) – Ct (IP)] was applied.

### Statistical analyses

Statistical analyses were performed with ANOVA, Student’s unpaired t-test using GraphPad Prism Ver. 9.2 (GraphPad La Jolla,CA) software, or as indicated in the legends. P-value is indicated by asterisks in the figures:*P≤0.05,**P<0.01,***P<0.001,****P<0.0001. Data from SVF cells were analyzed using ordinary one-way ANOVA with Holm-Šidák’s multiple comparisons test.

## Nonstandard abbreviations used

CD36: cluster of differentiation 36
HFD: high fat diet
IFRD: interferon related developmental regulator
KO: knockout
dKO: double knockout
RD: regular diet
SKMc15: skeletal muscle cDNA library clone15
SVF: stromal vascular fraction
TIS7: TPA Induced Sequence 7

## Acknowledgements

The authors thank Robert Kurzbauer for generation of dKO mice, Stephan Geley, Laura M. de Smalen, Laura De Gaetano and Karin Schluifer for the technical assistance, Christiane Heim for serum analyses and free fatty acid uptake measurements, Frans Stellaard for the analysis of fatty acids content in feces, Mayra Eduardoff for RNA processing for Affymetrix chip analysis, Alexander Magnutzki for advice with the statistical analyses of data, David Teis and Zlatko Trajanoski for critical reading of the manuscript. Furthermore, we would like to thank Dr. Paul Shore for providing us with the pGLCD36 construct. We are indebted to the staff at the Animal Facility of Innsbruck Medical University for their care of our mice. This work was supported by P18531-B12 and P22350-B12 grants from the Austrian FWF grant agency to Ilja Vietor and by the German Federal Ministry of Education and Research (Infrafrontier grant 01KX1012) to Martin Hrabe de Angelis.

## Author contributions

I.V., D.C., M.E., I.T., G.D., J.R., T.V. and L.A.H. designed the study. Experiments were performed by: Fig.1A DC, B VK+DC, C VK+GD, D,E GD, F DC; Fig.2A,B ThV, C IV, D,E,F,G ThV, H IV+ThV+KP; Fig.3A KP, B IV+ThV, C,D,E,F ThV+KP, G IV+TV, H,I IV+DH+ME+TV; Fig.4A IV+ME, B IV, C,D,E,F,G ThV, H ThV+KP, I ThV; Fig.5 KP; Fig.6 A,B IT+FS+DC, C,D,E KP; Fig.EV1A IV, B JR, C,D TV+ThV; Fig.EV2 TV+ThV; Fig.EV3A TV+ThV, B,C, D, E IV+TV; F TV+ThV; Suppl.Fig.1 A JR, B,C IT+PE+ED, D KP, E,F RG.

I.V., D.C., T.V. and L.A.H. wrote the manuscript.

**Figure EV1. Body length and respiratory exchange of dKO do not differ from WT mice. TIS7 and SKMc15 positively regulate adipocyte differentiation of MEF cells.**

A) Measurement of whole body length, including the tale of age-and sex-matched mice. n=14 and 17, respectively. Data shown are mean ± STD, *P<0.01

B) Indirect calorimetry measurements of WT and dKO animals. No significant differences were found in respiratory exchange ratios (RER) of WT and dKO mice.

C) Oil red O staining of MEFs following 8 days adipocyte differentiation protocol. Ectopic expression of TIS7 and/or SKMc15 induced adipocyte differentiation of TIS7 KO MEFs, size bar 20μm. Quantification of oil red O staining, n=3. Data shown are mean ± STD, ***P<0.001.

D) Ectopic expression of SKMc15 induced adipocyte differentiation of SKMc15 KO MEFs, size bar 20μm. Quantification of oil red O staining, n=3. Data shown are mean ± STD, ***P<0.001.

**Figure EV2. TIS7 and SKMc15 positively regulate adipocyte differentiation of MEF cells and inhibit DLK-1 expression. DLK-1 inhibits adipocyte differentiation of dKO MEFs.**

A) Ectopic expression of TIS7 and/or SKMc15 induced adipocyte differentiation of dKO MEFs, size bar 20μm. Quantification of oil red O staining, n=3, *** P<0.001. Ectopic expression of TIS7 and/or SKMc15 inhibited DLK-1 mRNA expression detected in dKO MEF cells. Normalized on GAPDH. Error bars indicate standard deviations, WT MEFs values were set as 1. Data shown are mean ± STD, ***P<0.001.

B) Cells from which the conditioned medium was transferred during the differentiation. Oil red O staining (panels on the left) after 8 days of adipocyte differentiation. Western blot detection of DLK-1 protein in differentiated cells.

C) Oil red O staining (left panel) of dKO control and shDLK-1 MEFs following 8 days adipocyte differentiation protocol. DLK-1 knockdown induced adipocyte differentiation of dKO MEFs, size bar 20μm. Western blot documenting (middle panel) the knockdown efficiency of two independent shRNA constructs targeting DLK-1. Quantification of oil red O staining, n=3. Data shown are mean ± STD, *** P<0.001.

**Figure EV3. DLK-1 regulates Hes1 expression, adipocyte differentiation of dKO MEF cells, and C/EBPα expression. dKO MEFs-secreted DLK-1 inhibits adipocyte differentiation of WT MEF cells.**

A) Hes1 mRNA expression detected in 8 days differentiated MEF cells. Normalized on GAPDH. Error bars indicate standard deviations, WT MEFs values were set as 1. Data shown are mean ± STD, ***P<0.001.

B) Oil red O staining (left panel) of 8 days adipocyte-differentiated WT control MEFs, WT MEFs treated with a recombinant DLK-1 or expressing ectopic DLK-1.

C) Quantification of oil red O staining shown in panel E, n=3. Data shown are mean ± STD, ***P<0.001.

D) DLK-1 mRNA expression detected in 8 days differentiated MEF cells.

E) C/EBPα mRNA expression detected in 8 days differentiated MEF cells. Normalized on GAPDH. Error bars indicate standard deviations, WT MEFs values were set as 1. Data shown are mean ± STD, ***P<0.001.

F) Oil red O staining of WT control MEFs (left image) following 8 days adipocyte differentiation protocol. WT cells differentiated in conditioned medium from dKO MEFs (middle image) and WT MEFs treated with conditioned medium from dKO MEFs expressing DLK-1 shRNA construct (right image). Size bar 20μm. Western blot documenting (middle panel) the knockdown efficiency of shRNA constructs targeting DLK-1. Quantification of oil red O staining, n=3. Data shown are mean ± STD, ***P<0.001.

**Supplementary figure 1. No difference between WT and dKO mice in food consumption. Expression patterns of TIS7 and SKMc15 in WT MEFs during adipocyte differentiation. SKMc15 knockout strategy.**

A) No significant genotype effect on food intake; when adjusted to the body mass, non-significantly increased in dKO mice (4.0 g WT and 3.4 g dKO mice). Data shown are mean ± STD.

B) Hepatic lipase serum concentrations of WT (n=4) and dKO (n=4) 11 weeks old male mice after 3 weeks of HFD. Data shown are mean ± STD.

C) Lipoprotein lipase serum concentrations of WT (n=4) and dKO (n=4) 8 weeks old male mice following 3 weeks of HFD. Data shown are mean ± STD.

D) TIS7 expression reaches its maximum on day 3 of MEF adipocyte differentiation. Data shown are mean ± STD. Student’s t-test P<0.01, n=3.

E) SKMc15 reaches its maximum expression on day 5 of MEF adipocyte differentiation. Data shown are mean ± STD. Student’s t-test P<0.05, n=3. All Ct values were normalized to GAPDH.

F) Modification of the SKMc15 gene locus and characterization of the targeted and null genotype. Homologous recombination with the targeting vector inserts one loxP site and a frt site flanked neomycin resistance gene (PGK–Neo) into intron 1 at position 105515 (AY162905). A second loxP site inserted behind the poly A tail into the 3’ UTR of the SKMc15 gene at position 102082 (AY162905). Further downstream were added 15 additional nucleotides, parts of intron 1, exon 2 (splice site: donor and acceptor) and the CDS of hrGFP (out of Vitality hrGFP Mammalian Expression Vector pIRES-hrGFP-2a; Stratagene). Arrows indicated the orientation of the hrGFP gene and the loxP sites. Southern blot analysis. The fragment detected by the Southern probe, was 9.6 kb in the WT (insert, band V) and 15 kb in the deleted locus (band IV).

G) PCR genotype analysis of SKMc15 targeted and deleted alleles. I: 426 bp WT SKMc15 PCR fragment. II: 1772 bp SKMc15 targeted allele PCR product. III: 1700 bp SKMc15 GFP screening PCR product can be amplified from the targeted (F) and the deleted allele (-/-), but not from the WT (+/+) locus.

H) TIS7 PCR detection in dKO mice. Left panel: detection of 1800 bp TIS7 WT allele; right panel: detection of 2100 bp TIS7 KO allele.

