## Supplementary figures and images for "The negative adipogenesis regulator DLK1 is transcriptionally regulated by TIS7 (IFRD1) and translationally by its orthologue SKMc15 (IFRD2)"

### Supplementary figures

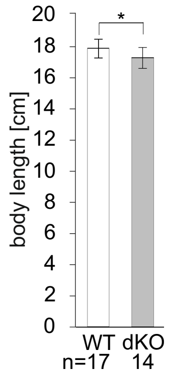

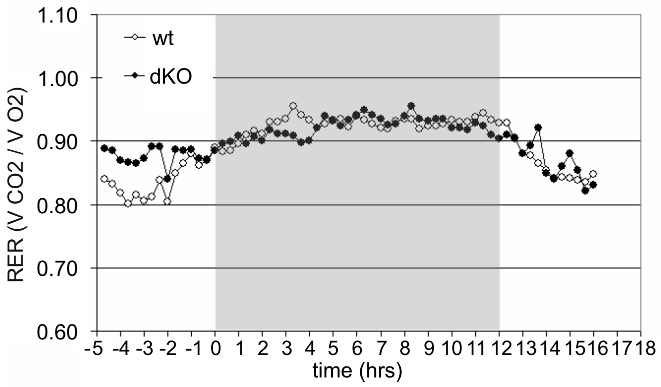

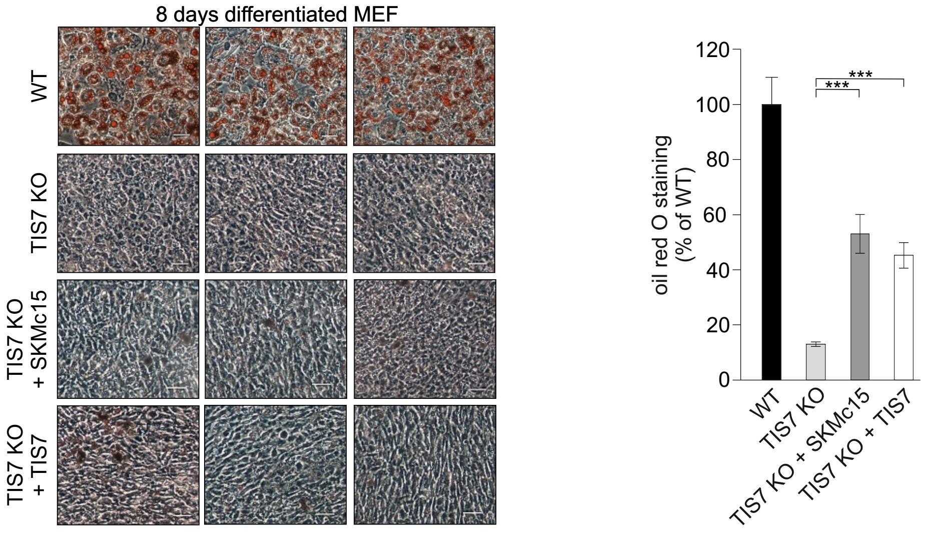

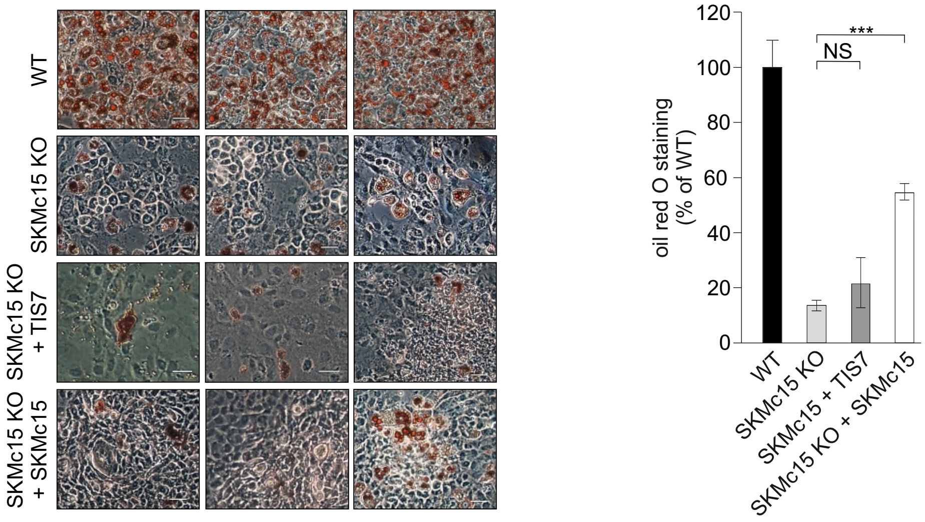

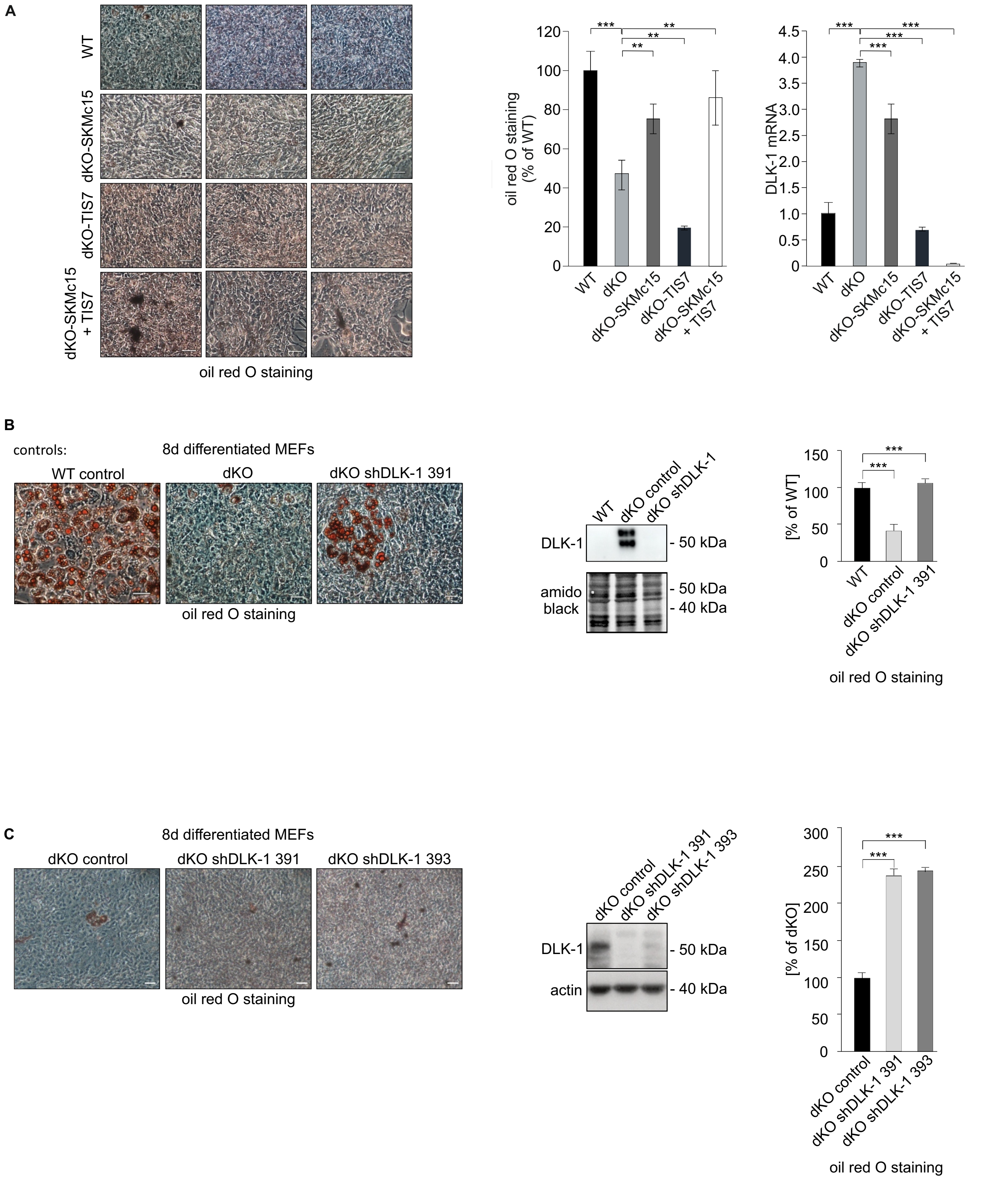

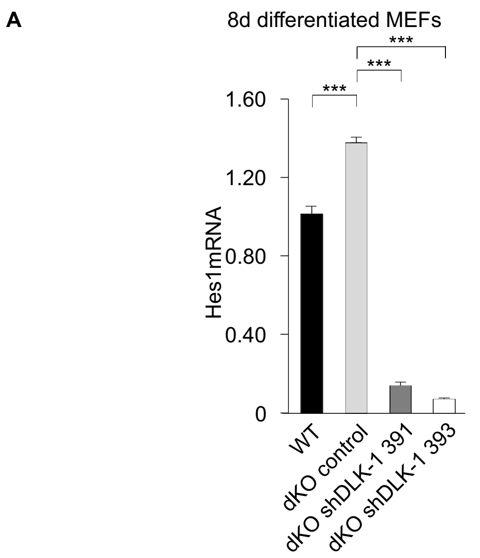

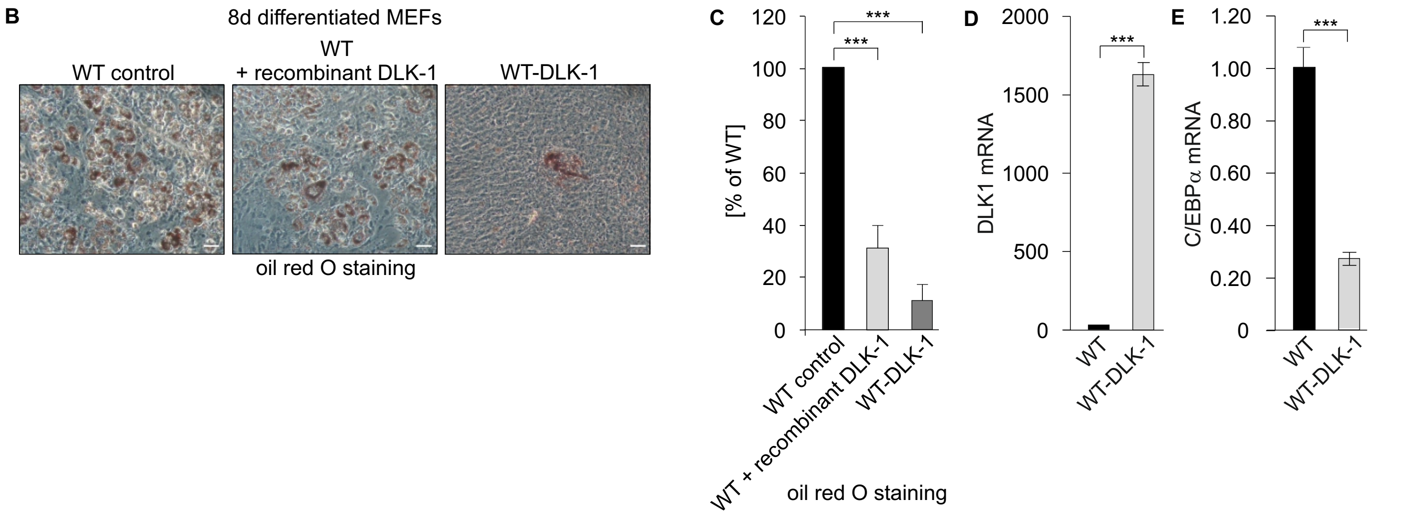

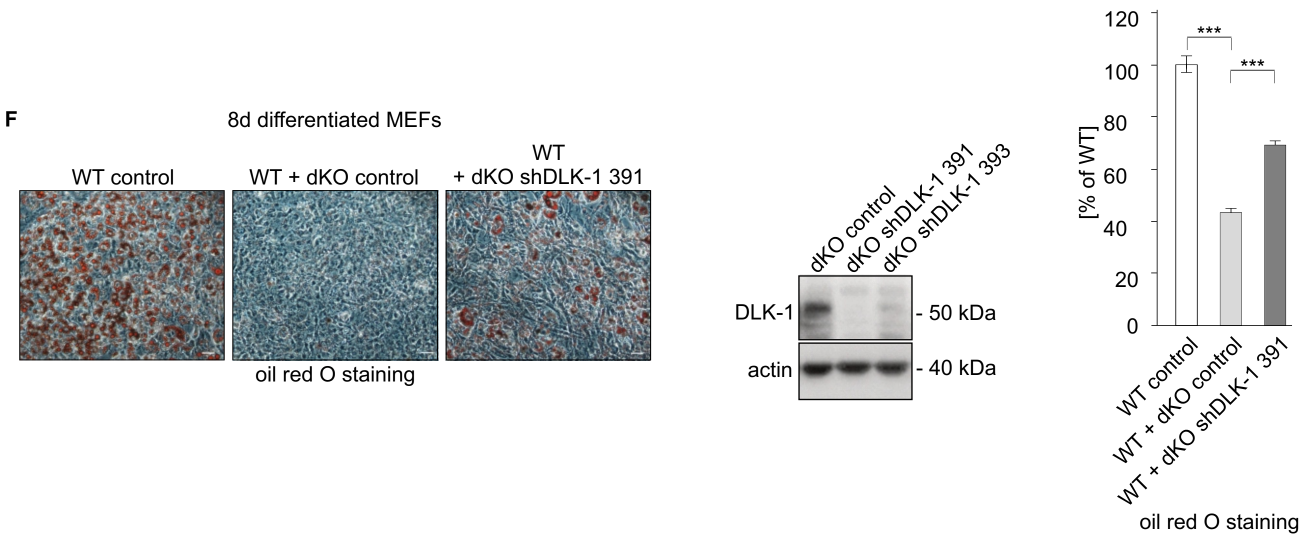

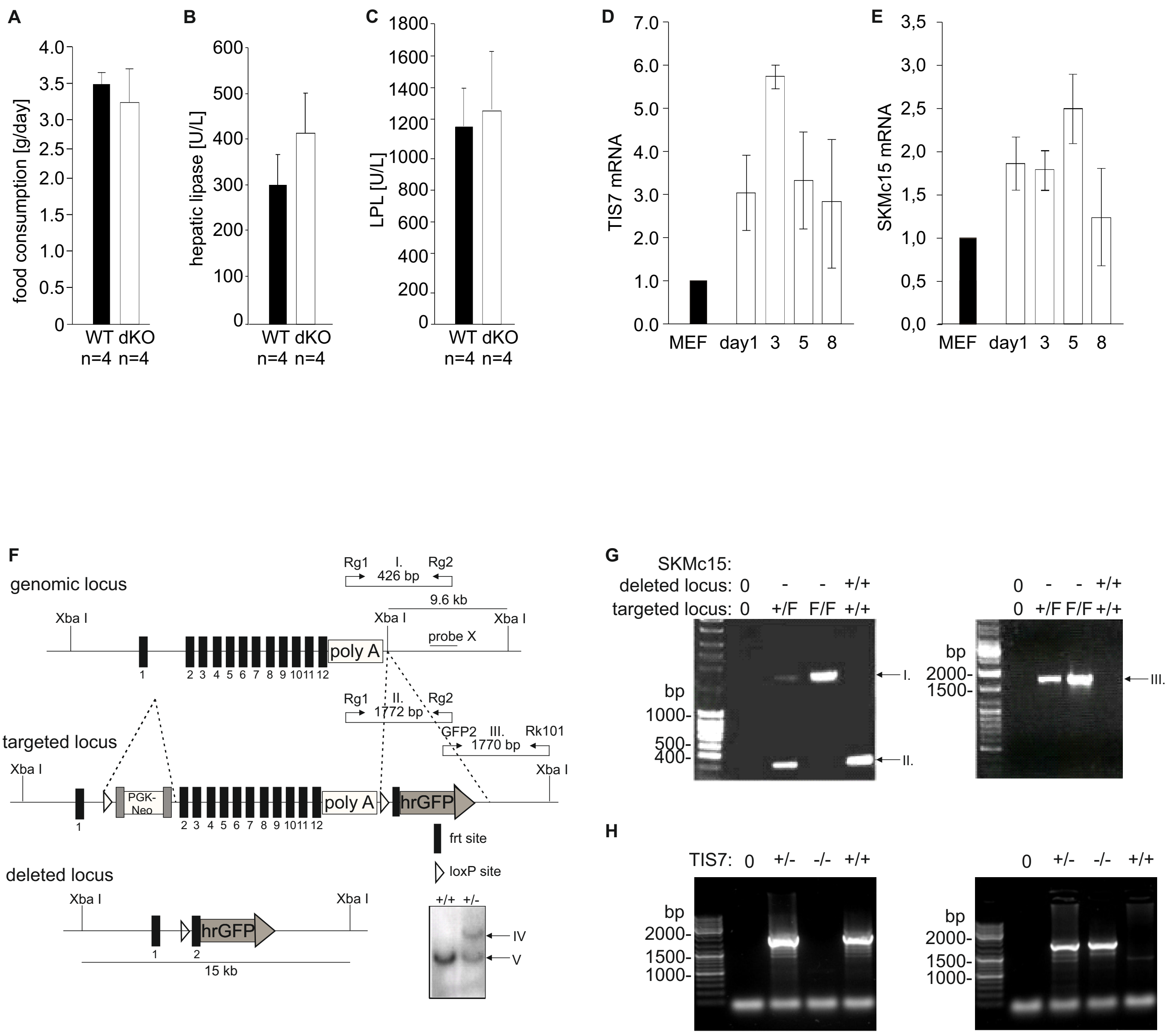
